# An activated wheat CC_G10_-NLR immune receptor forms an octameric resistosome

**DOI:** 10.1101/2025.08.26.672026

**Authors:** Guanghao Guo, He Zhao, Kaihong Bai, Qiuhong Wu, Lingli Dong, Lei Lu, Yongxing Chen, Yikun Hou, Jian Lu, Ping Lu, Miaomiao Li, Huaizhi Zhang, Gaojie Wang, Keyu Zhu, Baoge Huang, Xuejia Cui, Hongkui Fu, Chenchen Hu, Zhiying Chu, Xue Lyu, Sophien Kamoun, Chao Wang, Zhiyong Liu, Muniyandi Selvaraj, Jonathan D. G. Jones

## Abstract

Nucleotide-binding and leucine-rich repeat (LRR) receptors (NLRs) are widespread intracellular immune sensors across kingdoms. In plants, the G10-type coiled-coil (CC_G10_)-NLRs form a distinct phylogenetic clade that remains poorly characterized. Here, we identified a gain-of-function (GOF) mutant of *Wheat Autoimmunity 3* (*WAI3*), designated *WAI3^GOF^*, which encodes a constitutively activated CC_G10_-NLR protein due to an amino acid substitution. Cryo-EM structural analysis revealed that activated WAI3 assembles into a distinctive octameric resistosome. *Arabidopsis* RPS2, another CC_G10_-NLR, also forms an octamer, indicating a conserved structural property across monocot and dicot plants. The WAI3 resistosome mediates a prolonged and sustained increase in cytosolic calcium influx, facilitated by a unique channel architecture arising from its divergent CC domain configuration. Notably, this domain arrangement may be shared by many plant NLRs that lack the conserved EDVID motif in their CC domains. Our findings uncover a previously uncharacterized resistosome structure and provide insights into plant immune receptor plasticity.

## INTRODUCTION

Nucleotide-binding and leucine-rich repeat (NLR) proteins function as key immune receptors across diverse taxa (Jones et al., 2024). These intracellular proteins detect pathogen invasion and activate defense responses (Jones and Dangl, 2006). The NLR family is defined by its characteristic tripartite architecture: an N-terminal signaling domain that executes downstream immune activation, a regulatory nucleotide binding domain (NACHT in mammalians and NB-ARC in plants), and a C-terminal leucine-rich repeat (LRR) sensing domain (Jones et al., 2016). While the NB-ARC and LRR domains are structurally and functionally conserved in plants, the N-terminal signaling domains have diverged considerably, mediating immune signal output through distinct mechanisms (Huang et al., 2023).

Plant NLRs can be broadly categorized into two classes: Toll/interleukin-1 receptor/resistance protein- NLRs (TIR-NLRs) and coiled-coil NLRs (CC-NLRs) (Jubic et al., 2019). However, comprehensive phylogenomic analyses show extensive diversity within the CC-NLRs (Contreras et al., 2023; Lee et al., 2021; Seo et al., 2016). Recent cryo-electron microscopy (cryo-EM) studies have revealed that members from both types of NLRs assemble into oligomeric resistosomes that execute specific immune signalling functions upon pathogen effector detection (Förderer et al., 2022; Liu et al., 2024; Ma et al., 2020; Madhuprakash et al., 2024; Martin et al., 2020; Wang et al., 2019; Zhao et al., 2022). TIR-NLRs, such as Recognition of *Peronospora parasitica* 1 (RPP1) and Recognition of XopQ 1 (ROQ1), form tetrameric resistosomes upon effector recognition (Ma et al., 2020; Martin et al., 2020). This configuration brings together the N-terminal TIR domains to form two active sites that create NADase activity, producing nucleotide-derived small molecules that are perceived by downstream EDS1 (enhanced disease susceptibility 1) family heterodimers, ultimately triggering immune activation (Horsefield et al., 2019; Huang et al., 2025; Jia et al., 2022; Manik et al., 2022; Wan et al., 2019; Wu et al.; Xiao et al., 2025; Yu et al., 2022; Yu et al.). In contrast, CC-NLRs, including *Arabidopsis* HOPZ-ACTIVATED RESISTANCE 1 (ZAR1) and wheat stem rust resistance 35 (Sr35), assemble into pentameric structures upon activation. Their N-terminal CC domains form funnel-shaped structures that insert into the plasma membrane (PM) and function as calcium (Ca^2+^) permeable channels, triggering robust Ca²⁺ influx across the PM and elevating cytosolic Ca²⁺ ([Ca²⁺]_cyt_) levels, which is required and sufficient for inducing defense responses and cell death (Bi et al., 2021; Förderer et al., 2022; Wang et al., 2019). A specific group of CC-NLRs from asterid plants, termed NLRs required for cell death (NRCs), function downstream of other CC-NLRs for immune activation (Contreras et al., 2023). Two recent studies showed that these NRCs assemble into hexameric resistosomes, triggering immune responses by facilitating Ca^2+^ influx (Liu et al., 2024; Madhuprakash et al., 2024). The CC domain arrangement of NRCs resembles that of ZAR1 and Sr35, suggesting a conserved activation mechanism despite their different oligomeric states (Förderer et al., 2022; Liu et al., 2024; Madhuprakash et al., 2024; Wang et al., 2019).

An important feature shared by ZAR1, Sr35 and NRCs is the conserved EDVID motif in the α3 helix of their CC domains (Burdett et al., 2019; Förderer et al., 2022; Liu et al., 2024; Wang et al., 2019). Interaction of this motif with a conserved cluster of arginine residues in the LRR domain generates a unique “bent-back” CC domain conformation, contrasting with the TIR domain arrangement in activated TIR-NLR resistosomes (Förderer et al., 2022; Ma et al., 2020; Martin et al., 2020; Wang et al., 2019). The consequence of this CC domain arrangement is that while TIR domains sit on the same side as the NB-ARC domain in their respective resistosomes, the CC domains of activated ZAR1 and Sr35 are positioned opposite the NB-ARC domain. This unique CC–LRR interaction is believed to be stabilized in both the inactive and active conformations of CC-NLRs and may act to transduce signal from the LRR domain to the CC domain during activation (Burdett et al., 2019). The interaction is critical for their function, as mutations in either the EDVID motif or the arginine cluster abolish CC-NLR activity (Förderer et al., 2022; Liu et al., 2024). Intriguingly, a large fraction of CC-NLRs lack this motif in their CC domains, raising the fundamental question of how these EDVID-lacking CC-NLRs execute their function (Wroblewski et al., 2018).

Within the CC-NLR clade, two distinct CC-NLR phylogenetic sub-clades have been identified: RPW8- type CC-NLRs (CC_R_-NLRs) and G10-type CC-NLRs (CC_G10_-NLRs) (Contreras et al., 2023; Lee et al., 2021). The functions of CC_R_-NLRs, such as N requirement gene 1 (NRG1) and activated disease resistance 1 (ADR1), have been extensively studied. They serve as helper NLRs downstream of TIR signalling and detect conformational changes in EDS1 family heterodimers upon ligand binding, oligomerizing to form ion channels to mediate Ca^2+^ influx (Huang et al., 2025; Jacob et al., 2021; Xiao et al., 2025). CC_G10_-NLRs constitute another phylogenetically distinct subclade that includes the well-known resistance genes RESISTANCE TO PSEUDOMONAS SYRINGAE 2 and 5 (RPS2 and RPS5) (Day et al., 2005; Seo et al., 2016). These proteins are distinguished by the absence of the conserved EDVID motif in their CC domains (Lee et al., 2021). While significant advances have been made in understanding the molecular mechanisms of other NLR groups, the CC_G10_-NLRs remain largely uncharacterized, representing a critical knowledge gap in plant immune receptor biology.

In this study, we identified a gain-of-function mutant of *Wheat Autoimmunity 3* (*WAI3^GOF^*) in wheat, which displays autoimmune phenotypes including spontaneous hypersensitive response (HR)-like cell death lesions, constitutive immune activation, and enhanced resistance to wheat powdery mildew. *WAI3^GOF^* encodes an auto-active NLR belonging to the CC_G10_-NLR subgroup. We used transient expression and immunoprecipitation of a “funnel mutant” allele of WAI3^GOF^ in *Nicotiana* (*N.*) *benthamiana* to perform cryo-EM analysis and revealed that activated WAI3 assembles into an octameric resistosome–an unprecedented oligomeric state for plant NLRs. This octameric assembly appears to be a unique feature of the CC_G10_-NLR subgroup, as RPS2, another CC_G10_-NLR, also forms an octameric resistosome. Functional analysis in *N. benthamiana* demonstrates that WAI3^GOF^ induces a more prolonged and sustained increase in [Ca^2+^]_cyt_ compared to activated NRC4. Strikingly, structural analysis revealed that, unlike other reported CC-NLRs, the WAI3 resistosome features a unique CC domain positioning that mirrors the TIR domains in activated TIR-NLRs. This results in a previously undescribed resistosome architecture and an inverted membrane orientation. The unusual CC domain positioning likely stems from the absence of the conserved EDVID motif, which is known to mediate the CC–LRR interaction in other CC-NLR resistosomes, including ZAR1, Sr35, and NRCs (Burdett et al., 2019; Förderer et al., 2022; Liu et al., 2024; Wang et al., 2019). These findings uncover a previously uncharacterized assembly mechanism for the CC_G10_-NLR subgroup and fill an essential gap in our understanding of their immune signalling mechanisms.

## RESULTS

### WAI3^GOF^ triggers constitutive activation of defense responses in wheat

The M3405 mutant was identified from an ethyl methane sulfonate (EMS)-mutagenized population of the wheat line Zhongke331 (ZK331). At the adult plant stage, this mutant exhibited spontaneous HR-like cell death lesions, with F_1_ plants displaying intermediate phenotypes, indicating that the mutation is a semi-dominant, gain-of-function (GOF) allele (Figures, 1A and 1B). Genetic analysis of the M3405 × Liangxing 99 (LX99) progenies indicated that the HR-like cell death lesion phenotype was controlled by a single semi-dominant gene (Figure S1A). At the seedling stage, the mutant also exhibited HR-like cell death lesions and showed significantly enhanced resistance to powdery mildew (Figure 1C and Figure S1B). Trypan blue and 3,3’-diaminobenzidine (DAB) staining revealed extensive cell death and hydrogen peroxide (H_2_O_2_) accumulation associated with the lesions, respectively (Figures 1D and 1E). In contrast, wild-type plants showed no lesions, cell death, or H_2_O_2_ accumulation (Figures 1D and 1E). In addition, pathogenesis-related (*PR*) genes were significantly upregulated in leaves of the M3405 mutant compared to those of wild-type ZK331 plants (Figure 1F). Collectively, these results demonstrate that this GOF mutation constitutively activates immune responses, leading to an HR-like cell death. We therefore designated the gene as *Wheat Autoimmunity 3* (*WAI3*), following the naming convention established for two previously named autoimmune mutants in our lab, and named the GOF variant as *WAI3^GOF^*.

**Figure 1.**
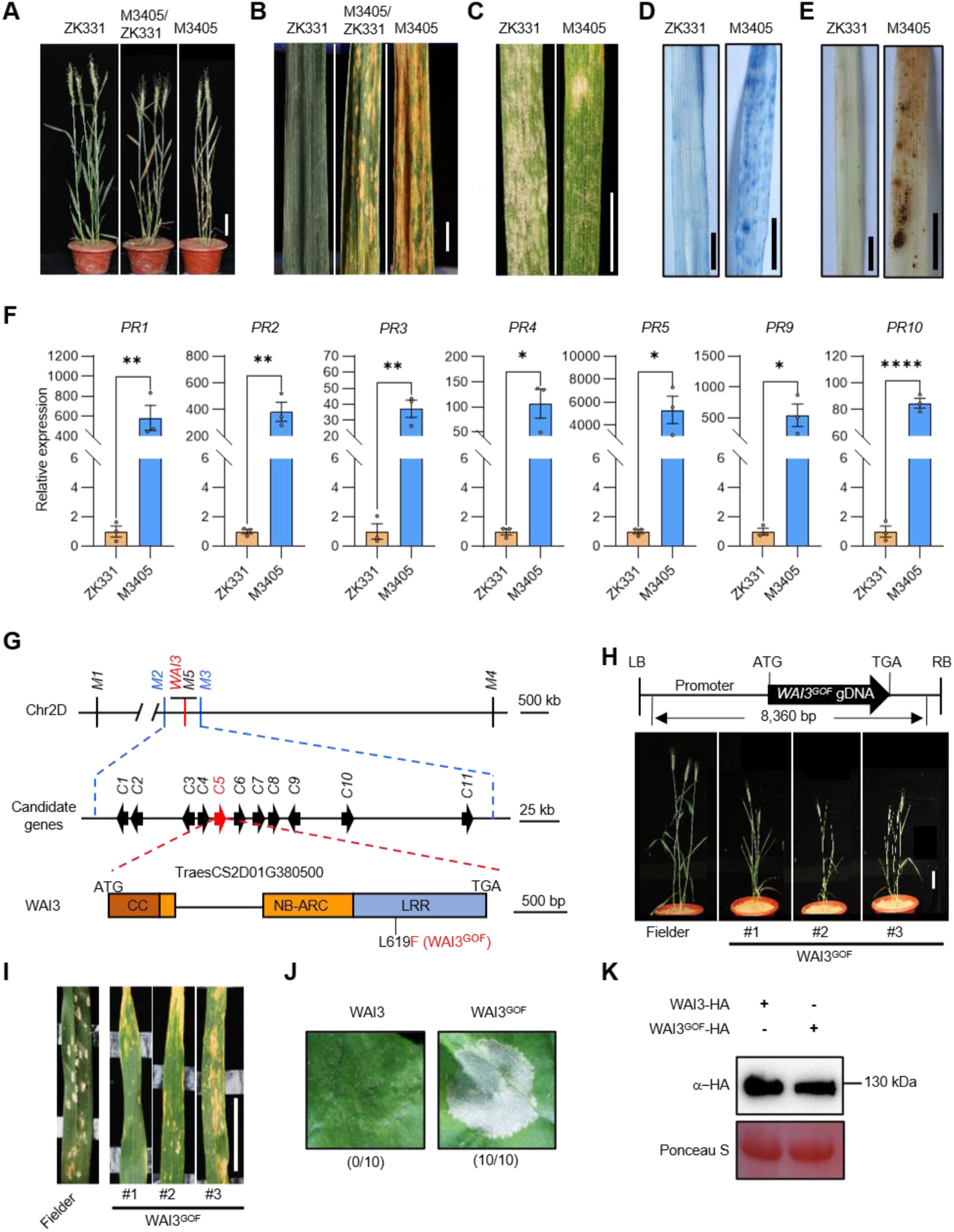
Identification of *WAI3* as the causal gene. (A and B) At the adult plant stage, M3405 plants exhibited spontaneous HR-like cell death lesions, with F_1_ plants displaying intermediate phenotypes, shown in whole plants (A; Scale bar, 10 cm) and leaf phenotype (B; Scale bar, 1 cm). (C) M3405 seedlings displayed significantly enhanced powdery mildew resistance relative to ZK331 at 14 days post-inoculation. Scale bar, 1 cm. (D) Trypan blue staining detected cell death in M3405 leaves but not in ZK331 leaves. Scale bar, 1 cm. (E) DAB staining revealed marked accumulation of H_2_O_2_ in M3405 leaves, but not in ZK331. Scale bar, 1 cm. (F) Pathogenesis-related (*PR*) genes were significantly upregulated in leaves of the M3405 mutant compared with those of ZK331 plants. Data are mean ± s.d. (n = 3, biologically independent samples). The wheat *ACTIN1* served as an internal control. Data were analyzed by two-tailed Student’s t test. Asterisks represent significant differences (\**P* < 0.05, \*\**P* < 0.01, \*\*\*\**P* < 0.0001). (G) Map-based cloning of *WAI3*. (H) Three independent T_1_ transgenic lines displayed an HR-like phenotype and no such phenotype was observed in the Fielder. Scale bar, 5 cm. (I) The transgenic plants showed significantly enhanced resistance to powdery mildew at the adult stage. (J) Transient expression of WAI3^GOF^ triggered a strong HR at 2 days post-infiltration (dpi), whereas expression of WAI3 did not. The representative figure is shown from 10 replicates and numbers below images indicate ratios of cell death leaves/all inoculated leaves. (K) Protein expression levels of WAI3 and WAI3^GOF^ in *N. benthamiana* leaves at 1 dpi. Rubisco staining with Ponceau S served as the loading control.

The *WAI3* locus was narrowed to a 600 kilobase (kb) region containing 11 high-confidence genes (designated *C1* to *C11*) on chromosome 2D (Figure 1G; Figure S1C and Table S1). Sanger sequencing revealed a mutation exclusively in the exon of gene *C5* (*TraesCS2D01G380500*) (Figure 1G). *C5* encodes an NLR protein comprising 919 amino acids (aa). In M3405, a C1855T nucleotide substitution resulted in a single amino acid substitution from leucine (Leu, CTT) to phenylalanine (Phe, TTT) at position 619 (L619F) in the LRR domain (Figure 1G and Figure S1D). The C1855T-derived marker *M5* showed co-segregation (Figure 1G). To confirm the candidate causal mutation, an 8,360-bp genomic fragment containing the *C5* gene with the C-to-T point mutation was transformed into the wheat cultivar Fielder, which harbors the wild-type *C5* gene, using *Agrobacterium*-mediated transformation. Three independent T_1_ transgenic plants exhibited HR-like cell death lesions resembling the M3405 mutant (Figure 1H) and exhibited significantly enhanced resistance to powdery mildew (Figure 1I), demonstrating that *C5* gene carrying the C1855T (L619F) mutation corresponds to *WAI3^GOF^*. To further validate the auto-activate nature of WAI3^GOF^, both WAI3^GOF^ and wild type WAI3 were transiently expressed in *N. benthamiana* leaves. Transient expression of WAI3^GOF^, but not wild-type WAI3 triggered a strong HR at 2 days post-infiltration (dpi) (Figures 1J and 1K). These findings demonstrate that the L619F mutation in the LRR domain is sufficient to induce the auto-activation of WAI3.

### Activated CC_G10_-NLR WAI3 forms an octameric resistosome

Following the identification of WAI3, a phylogenetic analysis was conducted to determine its evolutionary relationship with other plant NLRs. The result revealed that WAI3 belongs to the CC_G10_- NLR subclade, a distinct lineage that diverges significantly from other CC-NLRs (Figure 2A). This subgroup is characterized by the absence of the conserved EDVID motif in the CC domain (Seo et al., 2016; Wroblewski et al., 2018), and its functional mechanisms remain less well characterized compared to other CC-NLR and TIR-NLR groups. To elucidate the activation mechanism of this unique NLR subgroup, we sought to determine the structure of activated WAI3. We first examined whether WAI3^GOF^ forms higher-order complexes, similar to other activated NLRs, using blue native polyacrylamide gel electrophoresis (BN-PAGE). To circumvent cell death and facilitate protein expression, we generated a deletion variant of WAI3^GOF^ (WAI3^GOF/ΔN23^) by removing the α1-helix of the CC domain, the primary region responsible for cell death in other CC-NLR resistosomes (Förderer et al., 2022; Wang et al., 2019). The truncated WAI3^GOF/ΔN23^ variant exhibited robust expression while completely abolishing HR in *N. benthamiana* (Figures S1E and S1F). BN-PAGE revealed that WAI3^GOF/ΔN23^ was predominantly detected in slow-migrating forms at approximately 1,000 kDa, a pattern absent in WAI3, indicating oligomerization (Figure 2B). Noticeably, the observed size of the WAI3^GOF/ΔN23^ complexes (∼1000 kDa) significantly exceeds that expected for a pentamer (∼600 kDa) or hexamer (∼720 kDa), indicating that WAI3^GOF^ may assemble into a larger oligomeric structure (Figure 2B). To enable full-length protein production for structural analysis, we introduced residue substitutions in the N-terminal residues L12E/L15E (WAI3^GOF/L12E/L15E^), based on sequence homology with the ZAR1 and Sr35 resistosomes (Figure S1G). These mutations hamper the membrane association of the resistosome, thereby abolishing cell death without affecting oligomerization (Förderer et al., 2022; Liu et al., 2024). As expected, transiently expressed WAI3^GOF/L12E/L15E^ variant did not trigger HR in *N. benthamiana* (Figures S1H and 1I). We then co-expressed Flag- and Strep-tagged WAI3^GOF/L12E/L15E^ in *N. benthamiana* leaves and performed tandem purification to isolate oligomerized WAI3^GOF/L12E/L15E^ complexes. SDS-PAGE analysis revealed a single clean band corresponding to the expected size of WAI3^GOF/L12E/L15E^, confirming successful purification (Figure S2A). We then performed negative stain EM to further characterize the purified sample, which revealed octameric ring-like particles with high homogeneity (Figure 2C and Figure S2A). The observation of an octameric assembly aligns with the expected molecular weight of approximately 1,000 kDa observed in BN-PAGE (Figures 2B and 2C). Based on these results, we proceeded with cryo-EM analysis to determine the high-resolution structure of the WAI3 oligomeric complex.

**Figure 2.**
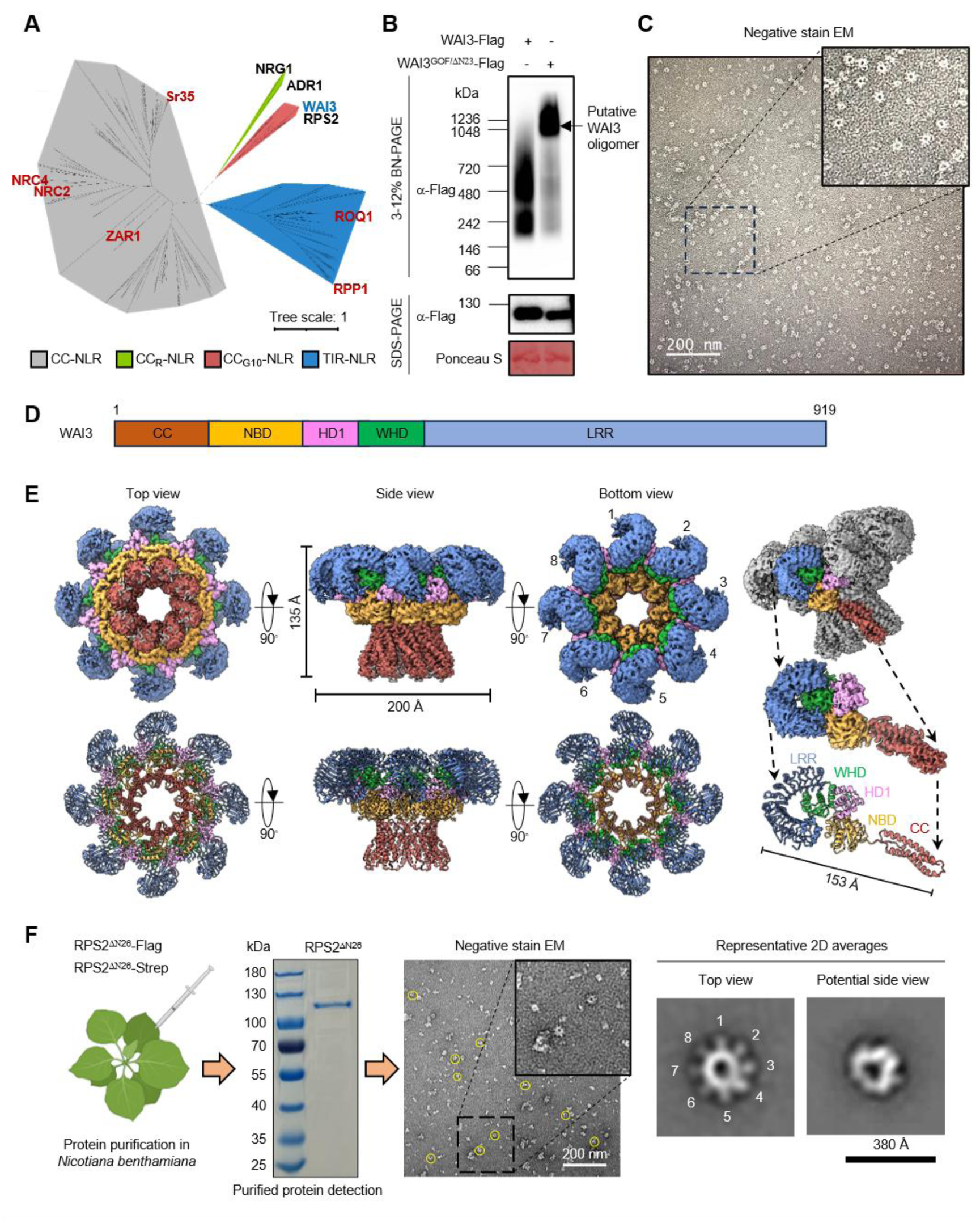
WAI3 belongs to the CC_G10_-NLR subgroup and forms an octameric resistosome upon activation. (A) Phylogenetic analysis positions WAI3 in the CC_G10_-NLR subgroup. Functionally characterized NLRs are labelled, with structurally resolved proteins shown in red and WAI3 highlighted in blue. (B) WAI3^GOF^ forms higher-order complexes compared with wild-type WAI3. An N-terminally truncated variant of WAI3^GOF^ (WAI3^GOF/ΔN23^) was used to reduce cell death and enhance protein expression. Proteins were transiently expressed in *N. benthamiana*, and total protein extracts were analyzed by BN- PAGE and SDS-PAGE, followed by immunoblotting with the indicated antibodies (shown at left). The experiment was independently repeated at least twice with consistent results. (C) Negative stain EM of purified WAI3^GOF/L12E/L15E^ reveals homogeneous, ring-like particles. This mutant was designed to disrupt plasma membrane localization and associated cell death, while retaining the N-terminal domain. Scale bar, 200 nm. (D) Schematic of the domain organization of WAI3. The same colour coding for domains is used throughout the study unless otherwise specified. (E) Cryo-EM density map and corresponding refined structural model of the octameric WAI3 resistosome, shown in top, side, and bottom views (left to right). (F) Expression and negative stain EM analysis of purified RPS2 protein. Representative 2D class averages show the octameric ring structures resembling the WAI3 resistosome.

A total of 7,043 cryo-EM images were collected and processed. Initial particle picking and reference-free 2D classification were performed on a subset of micrographs. The resulting 2D class averages revealed distinct resistosome particles in various orientations, displaying an octameric ring structure with cyclic symmetry (C8) and a narrow funnel-shaped end apparent when viewed from the side (Figures 2D and 2E; Figure S2 and Table 1). Additionally, two alternative configurations of the octameric resistosome were observed: one consisting of two octamers fused at their narrow funnel ends (designated as the N-to-N dual-octamer) and another with two octameric resistosome rings arranged back-to-back (designated as the B-to-B dual-octamer) (Figures 2D and 2E; Figure S2 and Table 1). The 2D class averages corresponding to these three distinct configurations were used as templates for particle picking across the entire dataset. Using these particles, we determined the 3D structures of the single octamer (C8 symmetry), the N-to-N dual-octamer (D8 symmetry), and the B-to-B dual-octamer (D8 symmetry) at overall resolutions of 3.95 Å, 4.5 Å, and 3.6 Å, respectively (Figures 2D and 2E; Figure S2 and Table 1). In all three reconstructions, the architecture of the single octameric resistosome was consistent with that observed within the N-to-N and B-to-B dual-octamers (Figure 2E and Figure S2). The final atomic model encompasses amino acid residues 36–919 of WAI3, excluding the regions 811–830 in the C-terminus and the first 35 residues in the N-terminus, as these segments were not resolved in the cryo-EM density map (Figure 2E and Figure S2). The α1-helix in the CC domain was not visible, likely due to its flexibility without membrane stabilization and this was similarly unresolved in Sr35, NRC2, and NRC4 resistosomes (Förderer et al., 2022; Liu et al., 2024; Madhuprakash et al., 2024).

**Table 1.**
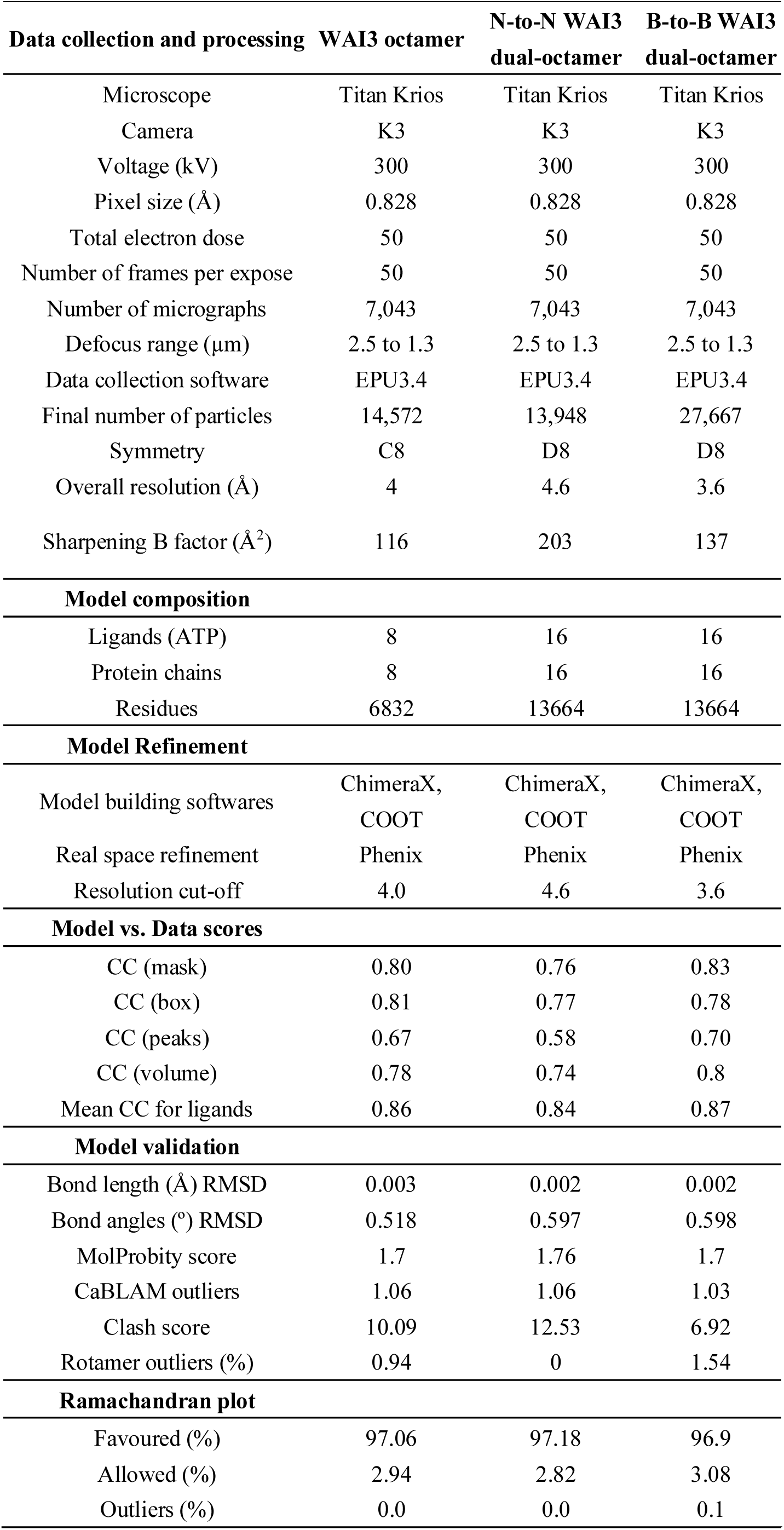
Statistics of the cryo-EM data collection, processing, and refinement.

While the N-to-N assembly have been repeatedly reported in Sr35 and NRC4 (Förderer et al., 2022; Liu et al., 2024), the B-to-B assembly was first identified in WAI3. The physiological significance of both these configurations remains unclear. Subsequent analyses focused exclusively on the single octamer assembly. The WAI3 octamer forms a ring-shaped assembly with a diameter of approximately 200 Å and a height of 135 Å. Similar to other subgroups of CC-NLRs, the active WAI3 structure comprises five distinct domains: a coiled-coil (CC) domain (residues 1–162), a nucleotide-binding domain (NBD) (residues 163–316), a helical domain 1 (HD1; residues 317–395), a winged-helix domain 1 (WHD; residues 396–495), and an LRR domain (residues 496–919), which contains 17 parallel β-strands (Figures 2D and 2E).

### The CC_G10_-NLR RPS2 also forms an octameric resistosome upon activation

Phylogenetic analysis suggested that the well-known resistance protein RPS2 belongs to the same CC_G10_- NLR subgroup as WAI3 (Figure 2A), prompting us to investigate its oligomerization state. In *Arabidopsis*, RPS2 activation is suppressed by its interacting partner RIN4 (Day et al., 2005). Transient expression of wild-type RPS2 without its suppressor triggers strong cell death in *N. benthamiana*, confirming the intrinsic activation nature of RPS2 (Figures S3A and S3B). To suppress cell death and enhance protein expression, we employed an N-terminal deletion variant (RPS2^ΔN26^) lacking the first 26 amino acids. This variant was confirmed to abolish HR cell death (Figures S3A and S3B). Co-expression of Flag- and Strep-tagged RPS2^ΔN26^ in *N. benthamiana* leaves enabled efficient tandem affinity purification with high purity (Figure 2F), which was subsequently analyzed by negative stain EM. We observed octameric ring assemblies, and potential side, and double-ring views (Figure 2F and Figure S3C). However, despite exhibiting a single band by Coomassie staining, the purified RPS2 sample showed high heterogeneity under EM, with only a minor fraction adopting oligomeric structures (Figure 2F and Figure S3C). Recently, RIN4 fragments were reported to act as potent cytosolic activators of RPS2 (Afzal et al., 2025), suggesting that transiently expressed RPS2 alone may not be completely activated. Nevertheless, these data clearly demonstrate that activated RPS2 can form octameric assemblies similar to the WAI3 resistosome (Figures 2E and 2F), indicating that the octamer formation might be a conserved property among CC_G10_-NLRs.

### Multiple interfaces stabilize the octameric WAI3 resistosome

Multiple interfaces, including CC, NBD, HD1, and WHD domains between adjacent protomers contribute to the stabilization of the WAI3 resistosome, while the LRR domain plays a minimal role (Figures 3A–3D). Along the structure of the resistosome, each protomer interacts closely with its neighboring protomer, burying a surface area of 4,980 Å² and forming a closed cyclic octamer with a large central solvent cavity. At a 4.5 Å cutoff for non-covalent interactions, 38 residues from one protomer interact with 35 residues of the adjacent protomer (Figure S4). At the interface between adjacent CC domains, the long N-terminal α2-helix (residues 38–69) from one protomer packs against the α3- helix (residues 75–105) and α4-helix (residues 110–136) from an adjacent protomer (Figures 3A and 3B). Among these multiple interactions, notable are hydrophobic interactions between A54 from one protomer with P75 and V79 from the neighboring protomer and an ionic bond formed by K47 with E128 of the neighboring protomer within the CC domains (Figure 3B). Within the NBD–NBD interface, E234, K255 and K260 forms a network of polar interactions with the adjacent protomer (Figure 3C). Additionally, the HD1 domain of one protomer engages the NBD and WHD domains of the neighboring protomer. In the HD1–NBD interface, F302, R308 and K328 make hydrophobic and polar interactions with the adjacent protomer (Figure 3D). For HD1–WHD interactions, E378 of HD1 forms a salt bridge with R457 in WHD domain of the adjacent protomer (Figure 3D). Notably, the CC–LRR interaction observed in ZAR1, Sr35, NRC2, and NRC4 resistosomes (Förderer et al., 2022; Liu et al., 2024; Madhuprakash et al., 2024; Wang et al., 2019) is absent in the WAI3 resistosome, which is described in detail in below. Furthermore, we find that D68 and K69 from the CC domain form a salt bridge and hydrophobic interactions, respectively, with the NBD domain of the neighboring protomer (Figures 3B and 3C). As previously reported (Förderer et al., 2022), the long linker (residues 138–163) between the CC and NBD domain also mediates oligomerization of the WAI3 resistosome.

**Figure 3.**
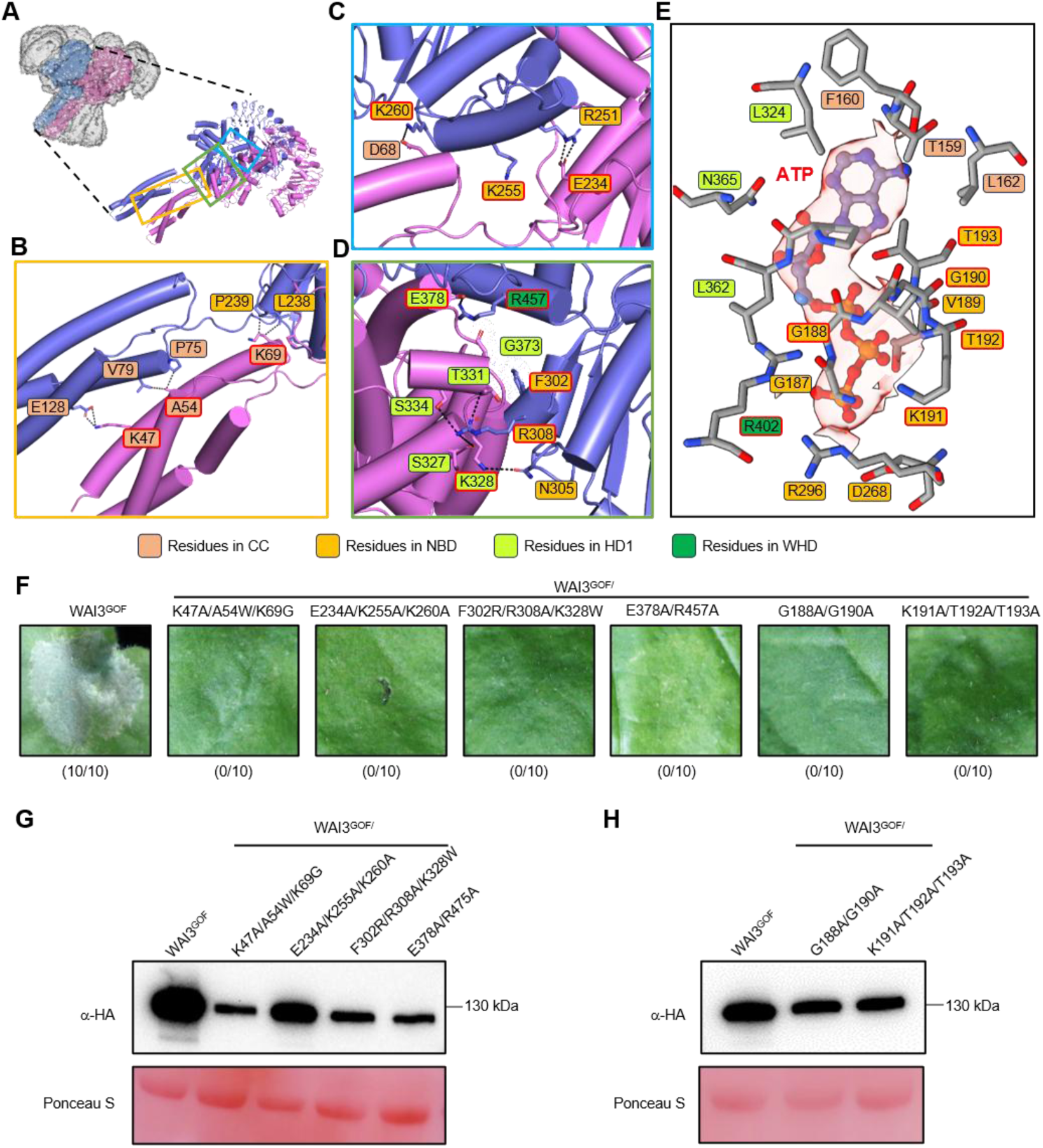
Multiple interfaces stabilize the octameric WAI3 resistosome. (A–D) Two adjacent protomers within the WAI3 resistosome, shown in different colours (A). Boxes in orange, blue and green indicate regions magnified in panels (B)–(D), respectively. (E) ATP was modeled into its corresponding density in the WAI3 octamer. Key residues surrounding the ATP within a 4.5 Å cut-off distance are labeled and shown as sticks. (F) Mutants targeting the residues at the interfaces shown in (B)–(D) abolished WAI3^GOF^-mediated cell death. Representative images from 10 biological replicates are shown, with the frequency of HR indicated below. (G and H) Protein expression levels of WAI3^GOF^ and the tested mutants in *N. benthamiana* leaves were assessed via SDS-PAGE and detected with an anti-HA antibody. Ponceau S staining of Rubisco served as a loading control.

To evaluate the contribution of these interactions to WAI3^GOF^-mediated HR, we conducted HR assays using targeted mutations at these interfaces. Four distinct groups of mutations targeting different protomer interfaces significantly reduced HR cell death (Figure 3F). Specifically, these groups were K47A/A54W/K69G at the CC–CC and CC–NBD interfaces; E234A/K255A/K260A at the CC–NBD and NBD–NBD interfaces; F302R/R308A/K328W at the NBD–HD1 interface; and E378A/R457A at the HD1–WHD interface. Both the WAI3^GOF^ and the tested mutants that reduced HR expressed well, indicating that the loss of cell death activity is not due to poor accumulation of these WAI3^GOF^ variants, confirming the importance of these interactions for resistosome assembly (Figure 3G).

ATP binding triggers a series of structural rearrangements that are crucial for NLRs to transition from their inactive to active states (Förderer et al., 2022; Liu et al., 2024; Wang et al., 2019). In the WAI3 resistosome, we observed a clear ATP density, with all interacting residues mapped within 4.5 Å. These residues span the NBD, HD1, and WHD domains of the NB-ARC region (Figure 3E). Mutations disrupting these interactions (G188A/G190A and K191A/T192A/T193A) markedly reduced the HR without significantly affecting protein stability (Figures 3F and 3H), highlighting the essential role of ATP binding in WAI3 function.

### The WAI3 resistosome triggers prolonged and sustained Ca^2+^ influx in planta

Increasing evidence demonstrates that oligomerized CC-NLRs (e.g. ZAR1, Sr35, NRC4) and CC_R_-NLRs (e.g. NRG1.1 and ADR1) form membrane-associated resistosomes with pore-forming capacity that mediate extracellular Ca^2+^ influx into the cytosol (Bi et al., 2021; Förderer et al., 2022; Jacob et al., 2021; Liu et al., 2024). To determine whether WAI3^GOF^ mediates Ca^2+^ influx, we transiently expressed WAI3^GOF^ and its mutant variants in stable transgenic *N. benthamiana* plants expressing the [Ca²⁺]_cyt_ reporter GCaMP3 (DeFalco et al., 2017). Fluorescence intensity was monitored over time to monitor Ca²⁺ dynamics.

Notably, compared to the empty vector (EV) and the control NRC4^DV^ (Liu et al., 2024), WAI3^GOF^ expression induces a prolonged and sustained elevation in [Ca²⁺]_cyt_ (Figures 4A and 4B). Two distinct phases of [Ca²⁺]_cyt_ increase were observed: an initial transient rise followed by a more prolonged and sustained phase, suggesting potential feed-forward regulation between Ca^2+^ signaling and WAI3 activity (Figures 4A and 4B). The [Ca^2+^]_cyt_ elevation differed between WAI3^GOF^ and NRC4^DV^, likely reflecting differences in their assembly mechanisms or protein stability in *N. benthamiana* leaves (Figures 4A and 4B). Application of LaCl₃, a known Ca²⁺ channel blocker, effectively abolished WAI3^GOF^-mediated Ca^2+^ influx (Figures 4A and 4B) and associated HR cell death (Figures 4C and 4D), demonstrating that WAI3^GOF^-mediated Ca^2+^ influx is dependent on channel activity and is sensitive to LaCl₃ inhibition. Structural and sequence analysis suggest that the L12E/L15E mutation impairs membrane binding outside the putative funnel region, while the D3K/E17K/D20K mutation likely disrupts the acidic inner lining of the α1-helix, which is presumed to be critical for Ca^2+^ permeability (Bi et al., 2021). In contrast to the robust [Ca²⁺]_cyt_ elevation mediated by WAI3^GOF^, neither of these membrane binding or the channel- disrupting mutants, induced significant [Ca^2+^]_cyt_ elevation or HR cell death (Figures 4E–4H; Figures S1H and S1I), further supporting the essential role of WAI3^GOF^ in mediating Ca^2+^ influx across the PM. Similar to WAI3^GOF^, activated RPS2 also mediates Ca²⁺ influx, suggesting that CC_G10_-NLRs share a common activation mechanism (Figures S3D–S3I).

**Figure 4.**
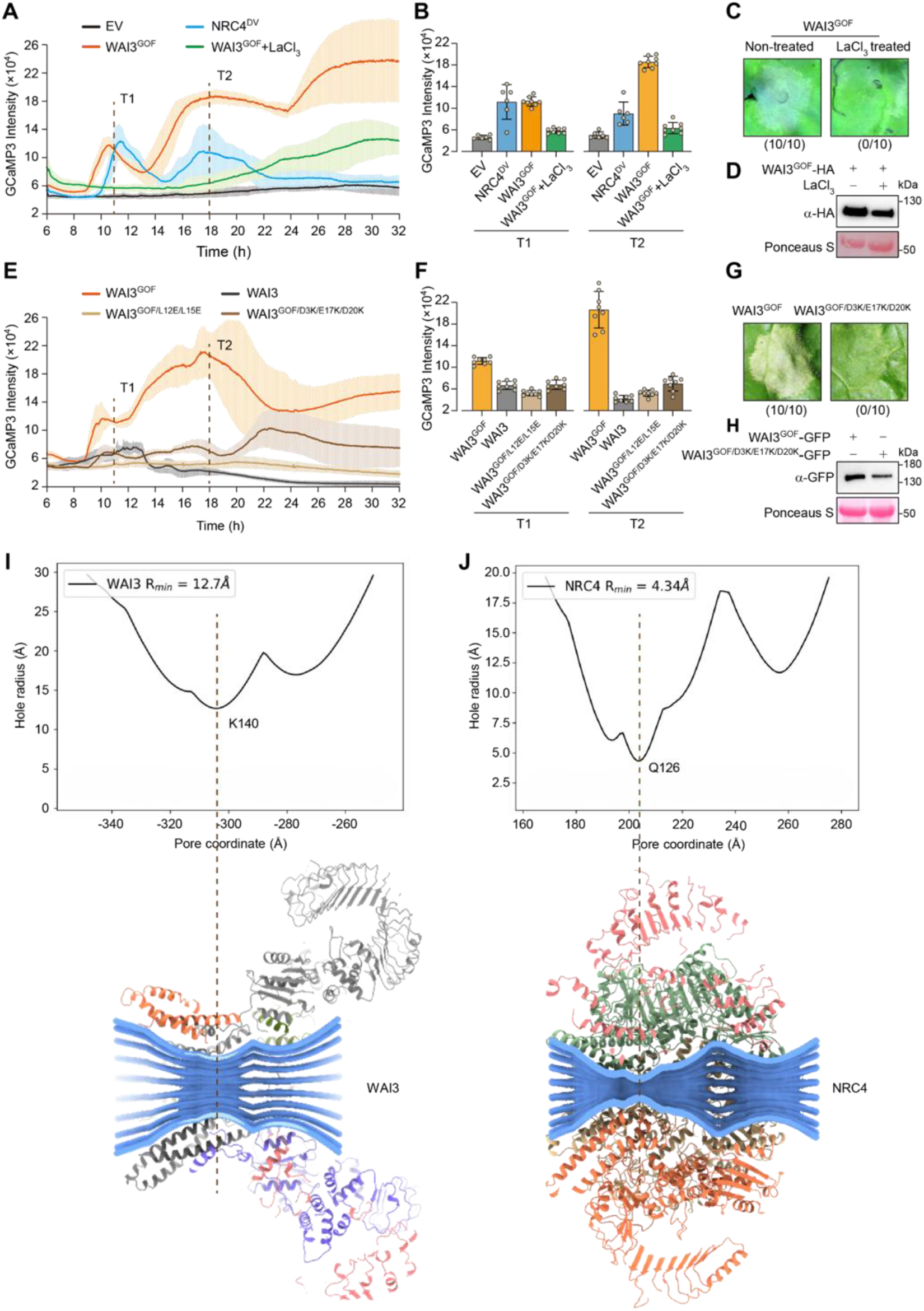
WAI3^GOF^ triggers unique [Ca²⁺]_cyt_ dynamics in *N. benthamiana* cells. (A) Expression of WAI3^GOF^ induces a robust and sustained Ca²^+^ influx in *N. benthamiana* cells, which is suppressed by 2 mM LaCl_3_ treatment. Time-course analysis of [Ca²^+^]_cyt_ dynamics was performed using *N. benthamiana* leaves expressing the Ca²^+^ reporter GCaMP3 (DeFalco et al., 2017). Leaves were infiltrated with *Agrobacterium* strains carrying the indicated constructs. GCaMP3 fluorescence intensity in leaf discs was measured over time to monitor [Ca²⁺]_cyt_ levels. The empty vector (EV) and auto-active NRC4^DV^ served as negative and positive controls. (B) Representative GCaMP3 fluorescence at two selected time points corresponding to the data in (A). (C and D) 2 mM LaCl_3_ suppressed WAI3^GOF^ -induced cell death in *N. benthamiana* (C) and protein expression was confirmed in (D). (E) Mutations at the predicted channel region disrupt WAI3^GOF^-mediated [Ca²^+^]_cyt_ elevation. WAI3^GOF/L12E/L15E^ represents mutations likely affecting membrane localization, while WAI3^GOF/D3K/E17K/D20K^ harbours substitutions predicted to impair ion conductance. (F) Representative GCaMP3 fluorescence at two selected time points corresponding to the data in (E). (G) The WAI3^GOF/D3K/E17K/D20K^ mutant abolishes WAI3^GOF^ -triggered cell death in *N. benthamiana*. Images were taken at 2 dpi. Numbers below images indicate ratios of cell death leaves/all inoculated leaves. (H) Immunoblot confirming protein expression in samples from (G). (I and J) Structural comparison of the channel regions of WAI3^GOF^ and NRC4^DV^ reveals distinct channel architectures. The ion permeation pathways of the activated WAI3 and NRC4 resistosomes, identified by PoreAnalyzer (Seiferth and Biggin, 2024), are depicted in blue. The residue at the narrowest point of each channel is highlighted with a dash line.

To further explore the relationship between [Ca²⁺]_cyt_ dynamics and WAI3^GOF^ resistosome assembly, we analysed the ion-conducting pathways formed by the WAI3^GOF^ octamer and the NRC4^DV^ hexamer using PoreAnalyzer (Seiferth and Biggin, 2024). Although the α1-helix of the CC domain was not resolved in neither structure, comparison of the pathways formed by the remaining structural elements revealed notable differences in pore architecture. The putative ion-conducting pore of the WAI3^GOF^ resistosome exhibits distinct architectural features compared to that of the NRC4^DV^ hexamer, particularly in overall geometry and minimum radius (Figures 4I and 4J). While the NRC4^DV^ hexamer pore narrows to a minimum radius of 4.3 Å at Gln126 (Figure 4J), the WAI3^GOF^ octamer pore maintains a notably wider minimum radius of 12.7 Å, with the primary constriction occurring at Lys140 (Figure 4I). This architectural difference may contribute to the distinct [Ca²⁺]_cyt_ dynamics observed in WAI3^GOF^ (Figures 4A and 4B). However, future studies, including experimentally resolved structures of the membrane-associated pore, will be essential to fully understand the functional relevance.

### The WAI3 resistosome reveals a unique molecular architecture

Compared to previously reported plant NLR structures, the WAI3 octamer represents the highest-order oligomerization observed among plant NLRs (Figure 5A). In addition to its considerable size, the WAI3 resistosome exhibits an unexpected domain organization compared to other structurally characterized CC-NLR resistosomes (Figures 5A and 5B). In previously reported CC-NLR resistosomes, the CC domains surrounding the central cavity and located on the same side as LRR domains. In contrast, the WAI3 resistosome features its CC domains next to NB-ARC domains, and opposite the LRR domains, with the three-helical bundle of the CC domain forming the stem of the funnel-shaped resistosome (Figures 5A and 5B). The distinct domain arrangements result in the different pore architectures and an alteration in the construction of walls of ion pathway described above (Figures 4I and 4J). Beyond structural differences in the pore architecture, the altered domain orientation in WAI3 results in a reversal of membrane topology with its LRR domains facing the cytosol, unlike in other CC-NLRs where the LRR domains face the membrane. The functional significance of this topological difference requires further investigation.

**Figure 5.**
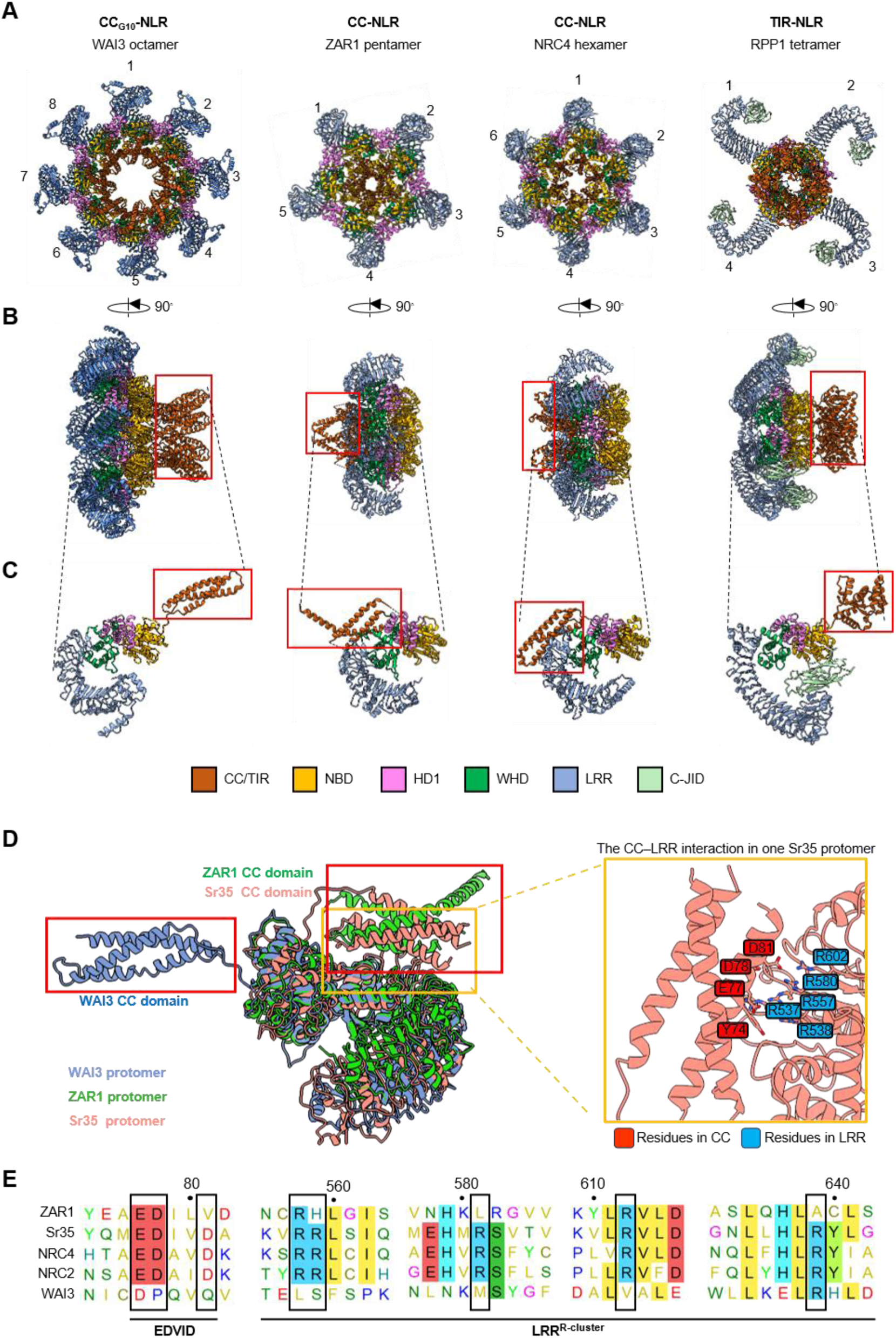
The WAI3 resistosome reveals unique molecular architecture. (A–C) Structural comparison of resistosomes assembled by NLRs from distinct phylogenetic groups reveals diverse oligomeric assemblies and domain organizations. Views shown: top (A), side (B), and protomer (C). Executor domains (CC or TIR) are highlighted in red boxes, illustrating distinct arrangements across NLR groups. PDB IDs: ZAR1 (PDB: 6J5T), NRC4 (PDB: 9CC8), RPP1 (PDB: 7DFV). (D) Compared to ZAR1 and Sr35, the WAI3 CC domain displays distinct structural organization (left), with the CC domain extended rather than bent back to contact the LRR domain. In Sr35, the EDVID motif mediates critical the CC–LRR interaction (right). (E) Sequence alignment of different CC-NLRs including ZAR1, Sr35, NRC2, NRC4, and WAI3, highlighting the presence or absence of the EDVID motif and the R-cluster. Conserved residues in the EDVID motif and arginine residues involved in the CC–LRR interaction in the Sr35 and NRC4 resistosomes are boxed.

We then investigated individual WAI3 protomers to determine the structural basis for its unique CC domain organization. The result revealed that, unlike protomers from other structurally characterized CC-NLRs, the CC domain in WAI3 is rotated approximately 180° relative to the hinge connecting it to the NBD domain (Figures 5C and 5D). In previously solved CC-NLR resistosomes (e.g. ZAR1, Sr35, NRC4, and NRC2), the CC domain helical bundle interacts with the LRR domain via contacts between the EDVID motif (in the CC domain) and an arginine-rich R-cluster (in the LRR domain) (Figure 5D) (Förderer et al., 2022; Liu et al., 2024; Madhuprakash et al., 2024; Wang et al., 2019). However, WAI3 lacks both the EDVID motif and the R-cluster, precluding such interactions (Figure 5E). The absence of this CC–LRR interaction likely contributes to the unique architecture of the WAI3 resistosome, including the inverted orientation of the NB-ARC and LRR domains compared to other CC-NLR resistosomes. Surprisingly, as a CC-NLR, the WAI3 resistosome configuration resembles TIR-NLRs (e.g., RPP1) more than previously characterized CC-NLRs, indicating possible different functional properties (Figures 5A–5C).

### CC-NLRs without the EDVID motif may exhibit similar CC domain organization

In the ZAR1, Sr35, NRC4 and NRC2 resistosomes, the CC domain directly interacts with the LRR domain via the conserved EDVID motif, which is essential for resistosome stability, signal transduction, and NLR function (Förderer et al., 2022; Liu et al., 2024; Madhuprakash et al., 2024; Wang et al., 2019). However, the EDVID motif is absent in a large proportion of CC-NLRs (e.g., being present in only 38% of *Arabidopsis* CC-NLRs), raising the key question of how EDVID-lacking CC-NLRs organize their domain architecture (Wroblewski et al., 2018). Based on CC domain sequence, *Arabidopsis* CC-NLRs have been classified into four groups. EDVID-lacking CC_R_-NLRs (e.g., NRG1) and CC_G10_-NLRs (e.g., RPS2) belong to groups A and B, respectively, while groups C and D include CC-NLRs such as ZAR1 and RPP8 (Wroblewski et al., 2018).

To explore whether different CC domain arrangements correlate with the presence or absence of the EDVID motif, we selected five representative CC-NLRs from each group, including three functionally characterized and two randomly selected uncharacterized members, and performed structural modelling using AlphaFold3 (Abramson et al., 2024). The results revealed that members from groups A and B (lacking the EDVID motif) adopt CC domain arrangements similar to WAI3, whereas those from groups C and D (with the EDVID motif) resemble the ZAR1-like architecture (Figure S5). Collectively, these findings indicate that the EDVID motif may serve as a critical determinant of CC domain organization, consistent with a recent independent study (Sulkowski et al., 2025). As the experimental structure of CC_R_-NLR NRG1 resistosome remains unresolved (Huang et al., 2025; Xiao et al., 2025), it remained unclear whether octameric assembly is associated with the absence of the EDVID motif.

## DISCUSSION

Although the structural architectures of multiple TIR-NLR and CC-NLR resistosomes have been elucidated (Förderer et al., 2022; Liu et al., 2024; Ma et al., 2020; Madhuprakash et al., 2024; Martin et al., 2020; Wang et al., 2019), the structural basis and nature of molecular assembly of CC_G10_-NLRs was unknown. Our structural investigation reveals that the CC_G10_-NLR WAI3 from the monocot crop plant wheat assembles into an octameric resistosome upon activation (Figure 2). An activated form of another CC_G10_-NLR, RPS2 from dicot plant *Arabidopsis thaliana*, exhibits similar octameric assemblies, indicating conserved oligomerization mechanism within this widely distributed phylogenetic clade. Notably, we found the CC domain configuration of WAI3 differs significantly from those of previously characterized pentameric and hexameric CC-NLR resistosomes (Figure 5). This structural divergence is likely attributed to the absence of the conserved EDVID motif in this CC_G10_-NLR clade–a feature critical for the CC–LRR interaction in other CC-NLR resistosomes. Interestingly, a substantial proportion of CC-NLRs also lack this motif, suggesting a universal, EDVID-independent assembly mechanism. Together, these findings establish CC_G10_-NLRs as a distinct class of resistosome with unique assembly principles and elucidate a conserved structural mechanism likely shared among EDVID-lacking CC-NLRs.

### Multi-phasic patterns of CC-NLR mediated Ca^2+^ influx

Previous studies have shown that CC-NLR resistosomes, such as ZAR1, Sr35 and NRC4, as well as CC_R_- NLR resistosomes like NRG1 and ADR1, induce Ca^2+^ influx across PM (Bi et al., 2021; Förderer et al., 2022; Jacob et al., 2021; Liu et al., 2024). Similarly, activated CC_G10_-NLR, including WAI3 and RPS2, also triggered robust Ca²⁺ influx (Figure 4A and Figure S3D), which requires octameric oligomerization on the PM to mediate Ca^2+^ permeability from the extracellular space (Figure 2B and Figures 4E–4H). The findings establish Ca^2+^ signaling as a convergence point in NLR activation and highlight its role as a critical second messenger during diverse NLR-mediated immunity processes. While earlier studies of ZAR1, Sr35, NRC4, NRG1, and ADR1 typically reported Ca²⁺ kinetic as a single transient peak, likely due to limited monitoring durations (Bi et al., 2021; Förderer et al., 2022; Jacob et al., 2021; Liu et al., 2024), our extended recording revealed a multi-phasic [Ca^2+^]_cyt_ elevation. This response includes an initial transient peak, consistent with previous reports, followed by a second, delayed phase of varying intensity across different CC-NLRs tested (Figure 4A and Figure S3D), suggesting a potentially conserved mechanism. These tissue-level Ca^2+^ oscillations resemble single-cell Ca^2+^ oscillations (Charpentier et al., 2016), and are likely regulated by feedback and/or feed-forward loops that influence cell survival and/or death, opening new avenues for future investigation.

### Different oligomeric states of plant resistosomes

Plant NLRs appear to adopt a range of oligomeric states, each characterized by a distinct protomer number. In resistosome structures, the HD1 and WHD domains constitute the resistosome rim, connected by a helix and a hinge region (Figure S6A). Structure superposition reveals variable inter-domain movement between HD1 and WHD domains (Figure S6B). The angle between HD1 and WHD domains was quantified based on 3 conserved hydrophobic residues (e.g., WAI3 I456, L410 and W379) in pentameric, hexameric and octameric assemblies. Experimental structures exhibit different angles: WAI3 octamer (80°), NRC2 hexamers (88°) and ZAR1 pentamer (92°) (Figure S6C). Notably, AlphaFold (AF)- predicted WAI3 (82°) closely matched its experimental octameric angle (80°). This correlation suggests that the HD1–WHD angle, derived from AF-predicted monomer models, can in principle predict NLR oligomeric state. Consistent with this, AF-modelled RPS2 (84°, near WAI3) suggested an octamer, matching the negative stain data (Figure 2F). In addition, AF-modeled NRG1 (94°, near ZAR1) predicted a pentamer, aligning with cryo-EM data of the EDS1-SAG101-NRG1^ΔN57^ complex where a minor fraction resembled pentameric particles (Huang et al., 2025). As more experimental structures become available, this angle can be further refined to deduce the oligomeric state of AF-modelled CC-NLRs.

A conserved C-terminal amphipathic helix within WHD (e.g., WAI3 479–491) facilitates the HD1–WHD domain movement. Hydrophobic residues projecting from one face of this helix (e.g., WAI3 V480, I481, L484, W487, V489) engage a complementary hydrophobic pocket (e.g., WAI3 V396, F400, L403, F407, L419) (Figure S6D). This interaction facilitates domain movement during activation from resting state of NLRs, effectively “greasing” the required hinge motion analogous to mechanisms in membrane transporters (Wöhlert et al., 2015). This amphipathic pivot helix is structurally conserved in ZAR1, NRC2, and RPP1 resistosomes (Figure S6E). We propose that the precise HD1–WHD angle dictates the rim unit geometry, thereby determining the number of monomers that can assemble into a closed symmetric ring. Variation in the side chain size, hydrophobicity and packing density of atoms of residues within this pivotal interface can directly modulate the achievable hinge angle range and, consequently, the oligomeric state.

### Dual-octamer formation

Beyond the octameric resistosome, we identified two additional dual-octamer states of WAI3 (Figure S2). In one arrangement, two octamers dock through their LRR domains, forming a B-to-B configuration. In the other, individual octamers dock via the tip region of their N-terminal CC domains, resulting in an N-to-N arrangement (Figure S2). These higher-order assemblies introduce dihedral symmetry, in addition to the intrinsic 8-fold cyclic symmetry of the octameric resistosome. Consequently, the structural analysis of the WAI3 resistosome reveals three distinct protein-protein interaction interfaces: (i) inter-protomer contacts within the octameric assembly, (ii) back-to-back interactions mediated by the LRR domains in the B-to-B dual octamer, and (iii) CC-domain-mediated interactions in the N-to-N dual-octamer (Figure S2). Notably, assemblies resembling the N-to-N arrangement have also been observed in the Sr35 and NRC4 resistosomes (Förderer et al., 2022; Liu et al., 2024). While the observed B-to-B arrangement has not been previously reported, it remains unclear whether this assembly is a result of overexpression artifacts. However, it is plausible that the oligomerization mediated by LRR domains in this case, may reflect a potential role of individual LRR domain of WAI3 to serve as a platform for protein-protein interactions with regulatory or downstream signaling components in vivo.

## ACKNOWLEDGMENTS

We thank Natasha Lukoyanova (Birkbeck, University of London, London, UK) for support with cryo-EM imaging and revising the manuscript. We thank the bioinformatics team of TSL and Structural biology platform of JIC for efficient support in data transfer and storage and computing access. We thank Jake Richardson for his excellent maintenance of the JIC-electron microscopy facility. We thank Mark Youles, Liam Egan for providing the Golden Gate modules for cloning. We thank AmirAli Toghani for helpful discussion. We thank all members of the TSL and IGDB Support Services for their invaluable assistance. GCaMP3 *N. benthamiana* line was kindly provided by Keiko Yoshioka. This study was supported by the National Key Research and Development Program of China (2022YFF1002800), the National Natural Science Foundation of China (U21A20224), Key Research and Development Program of Zhejiang (2024SSYS0099). Cryo-EM data for this investigation were collected at the Birkbeck ISMBEM facility, which was supported from the Wellcome Trust (202679/Z/16/Z and 206166/Z/17/Z).

G.G. was supported by the special project of long-term overseas training for young scientific and technological talents in 2022 of CAS.

## AUTHOR CONTRIBUTIONS

Conceptualization: G.G., Z.H., C.W., Z.L., M.S., J.D.G.J; Methodology: G.G., Z.H., C.W., M.S, N.L., X.L; Investigation: G.G., Z.H., M.S, K.B., Q.W., L.D., L.L., Y.C., Y.H., J.L., P.L., M.L., H.Z, G.W., K.Z., B.H., X.C., H.F., C.H., Z.C., X.L.; Visualization: M.S., Z.H, G.G.; Supervision: C.W., S.K., Z.L., J.D.G.J, Writing–original draft: G.G., Z.H., M.S., K.B.; Writing–review & editing: G.G., Z.H., M.S, K.B., S.K., C.W., Z.L., J.D.G.J.

---

**Figure S1.**
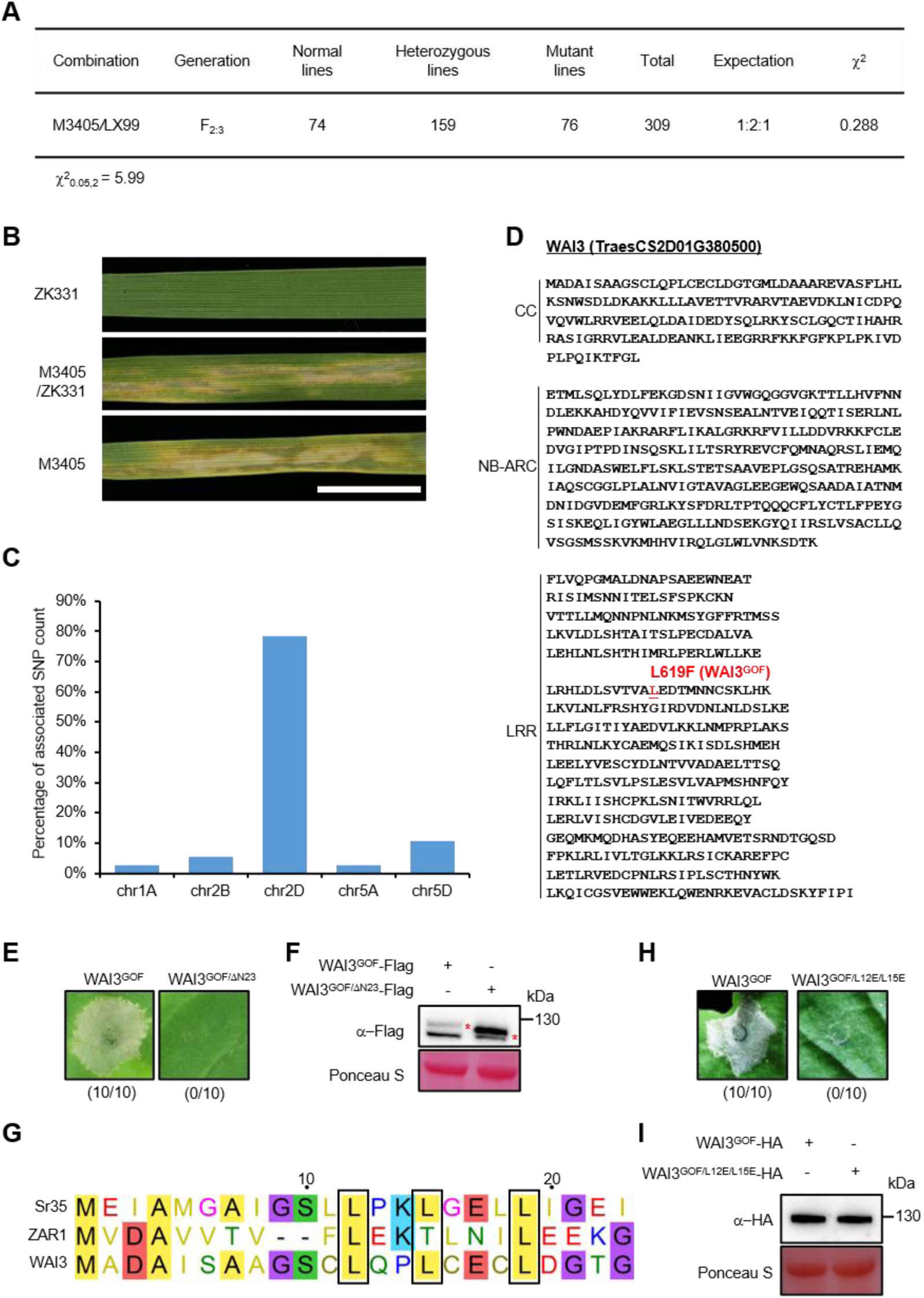
WAI3^GOF^ is a gain-of-function CC-NLR that requires an intact CC domain for function. (A) Genetic analysis of the M3405 × LX99 progenies indicated that the HR-like lesion phenotype was controlled by a single semi-dominant gene. (B) At the seedling stage, M3405 plants exhibited spontaneous HR-like cell death lesions, with F_1_ plants displaying intermediate phenotypes. Scale bar, 1 cm. (C) Single nucleotide polymorphisms (SNPs) associated with WAI3 were mapped to chromosome 2D in wheat using BSR-seq analysis. (D) Domain analysis of WAI3, with the gain-of-function (GOF) mutation in WAI3^GOF^ highlighted in red. (E) The deletion variant WAI3^GOF/ΔN23^, lacking the α1-helix, completely abolished WAI3^GOF^-mediated HR in *N. benthamiana* leaves at 2 dpi. Numbers below each image indicate the ratio of leaves showing cell death to the total number of inoculated leaves. (F) Protein expression levels of WAI3^GOF^ and WAI3^GOF/Δμ23^ in *N. benthamiana* leaves at 1 dpi. Rubisco staining with Ponceau S served as the loading control. (G) Protein sequence alignment highlights the conserved leucine (L) residues in the α1-helix, indicated by black boxes. (H) The L12E/L15E mutant (WAI3^GOF/L12E/L15E^) markedly suppressed WAI3^GOF^-mediated HR in *N. benthamiana* leaves at 2 dpi. Numbers below each image indicate the ratio of leaves showing cell death to the total number of inoculated leaves. (I) Protein expression levels of WAI3^GOF^ and WAI3^GOF/L12E/L15E^ in *N. benthamiana* leaves at 1 dpi. Rubisco staining with Ponceau S served as the loading control.

**Figure S2.**
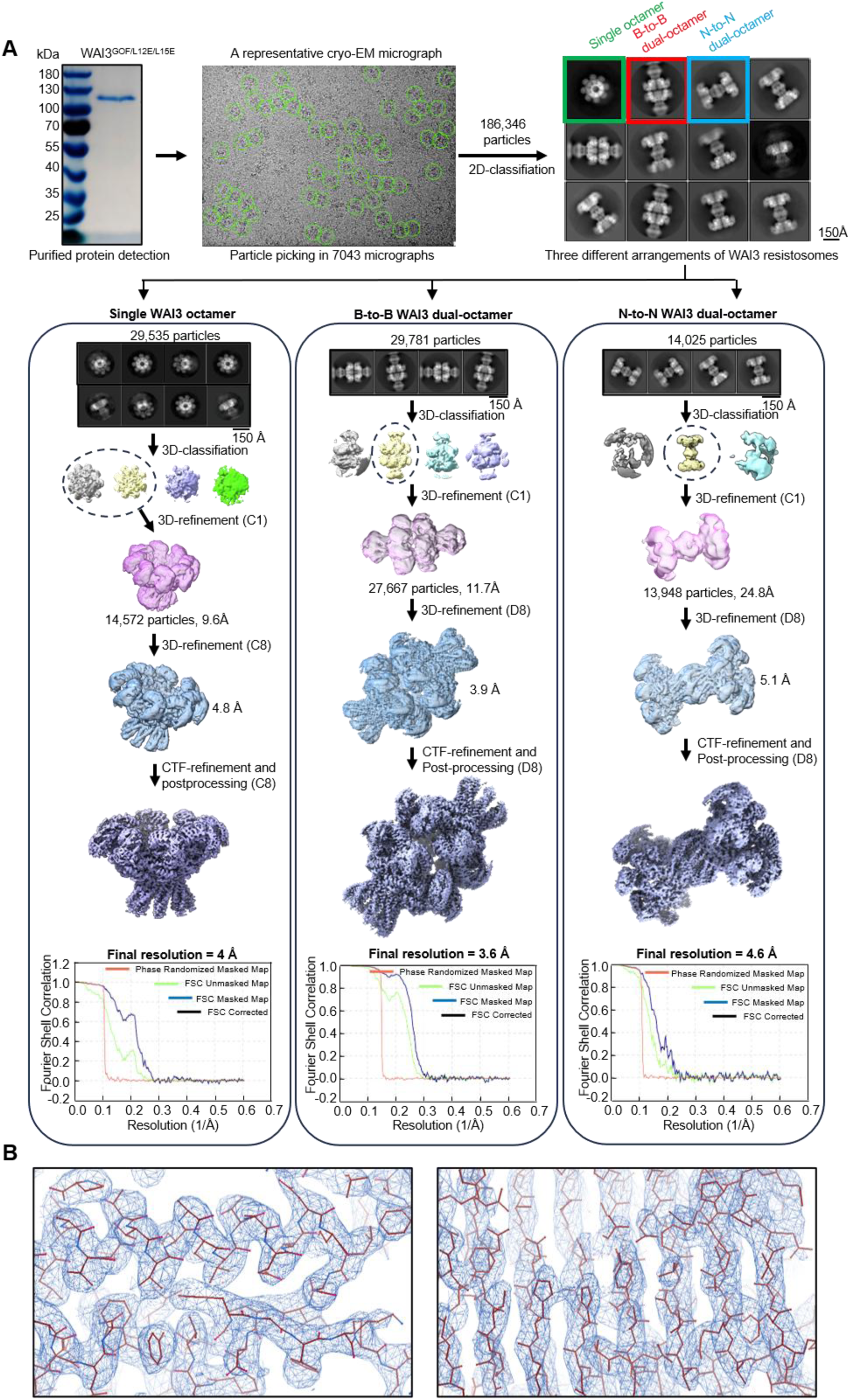
Flowchart of cryo-EM data processing and 3D-restruction and representative views of cryo-EM map density. (A) Data processing workflow, from purified protein samples to the final 3D reconstruction of WAI3 resistosomes. The Fourier Shell Correlation (FSC) curve of the final density map is shown to indicate resolution estimation. (B) The representative cryo-EM map densities of the WAI3 resistosome.

**Figure S3.**
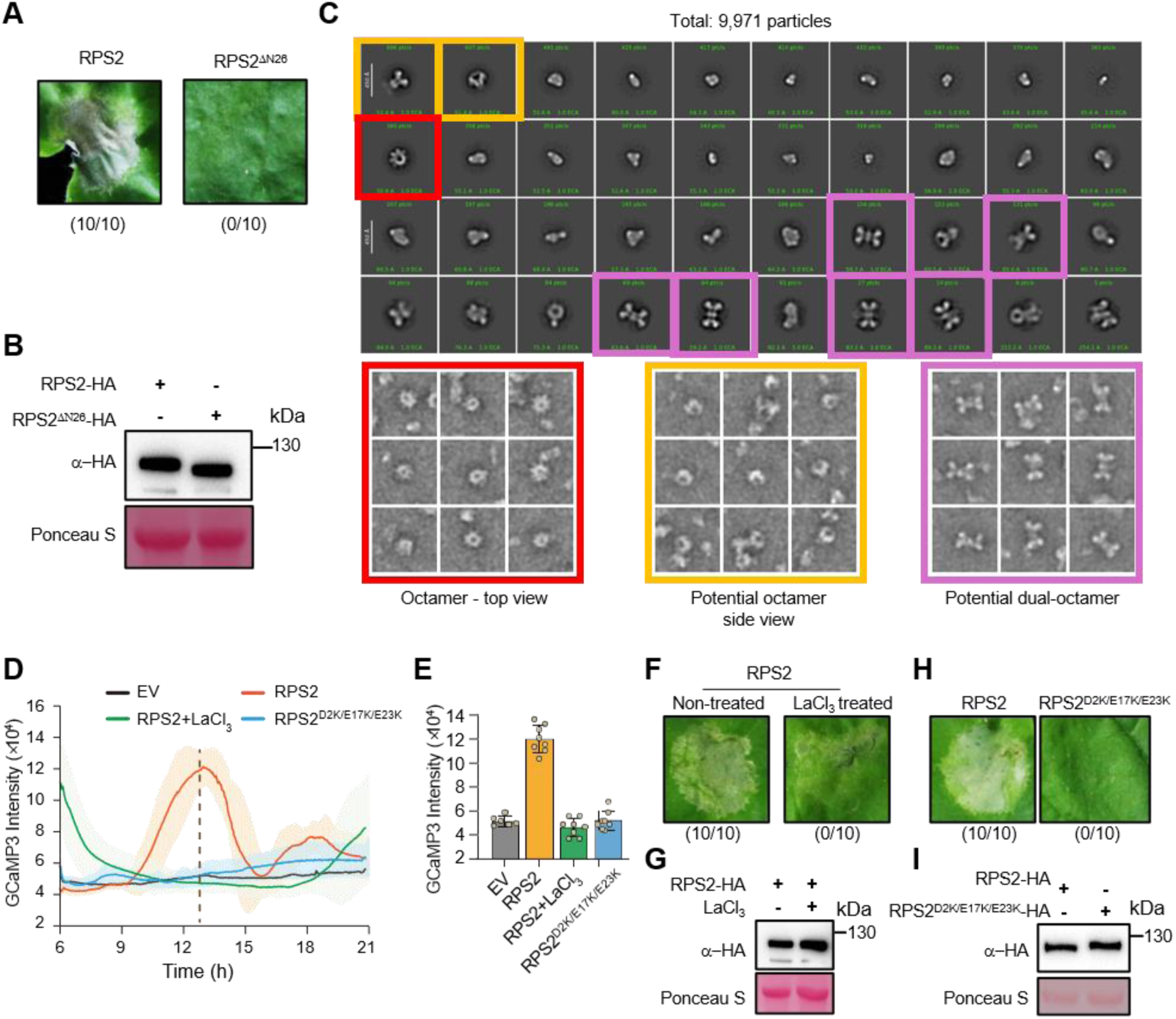
Negative stain EM of RPS2, and [Ca^2+^]_cyt_ dynamics triggered by RPS2 expression. (A) The N-terminal α1-helix deletion mutant (RPS2^ΔN26^) abolish RPS2-mediated HR phenotype in *N. benthamiana* leaves at 2 dpi. Numbers below images indicate ratios of cell death leaves/all inoculated leaves. (B) Protein expression levels of RPS2 and RPS2^ΔN26^ in *N. benthamiana* leaves at 1 dpi. Rubisco staining with Ponceau S served as the loading control. (C) 2D classification of negatively stained RPS2 particles. Top: 2D class averages showing RPS2 octamer (top view), potential side view, and dual octamer. Bottom: corresponding individual particles from raw images. (D) Expression of wild-type RPS2 induces a robust [Ca^2+^]_cyt_ influx in *N. benthamiana* cells, which is inhibited by treatment with 2 mM LaCl₃. In contrast, the RPS2^D2K/E17K/E23K^ channel mutant fails to trigger Ca^2+^ influx. (E) Representative GCaMP3 fluorescence at the selected time point corresponding to the data in (D). (F and G) RPS2-induced HR in *N. benthamiana* was suppressed by 2 mM LaCl_3_ (F). Protein expression was confirmed in (G). (H and I) RPS2^D2K/E17K/E23K^ abolished HR induced by RPS2 (H) and protein expression was confirmed in (I).

**Figure S4.**
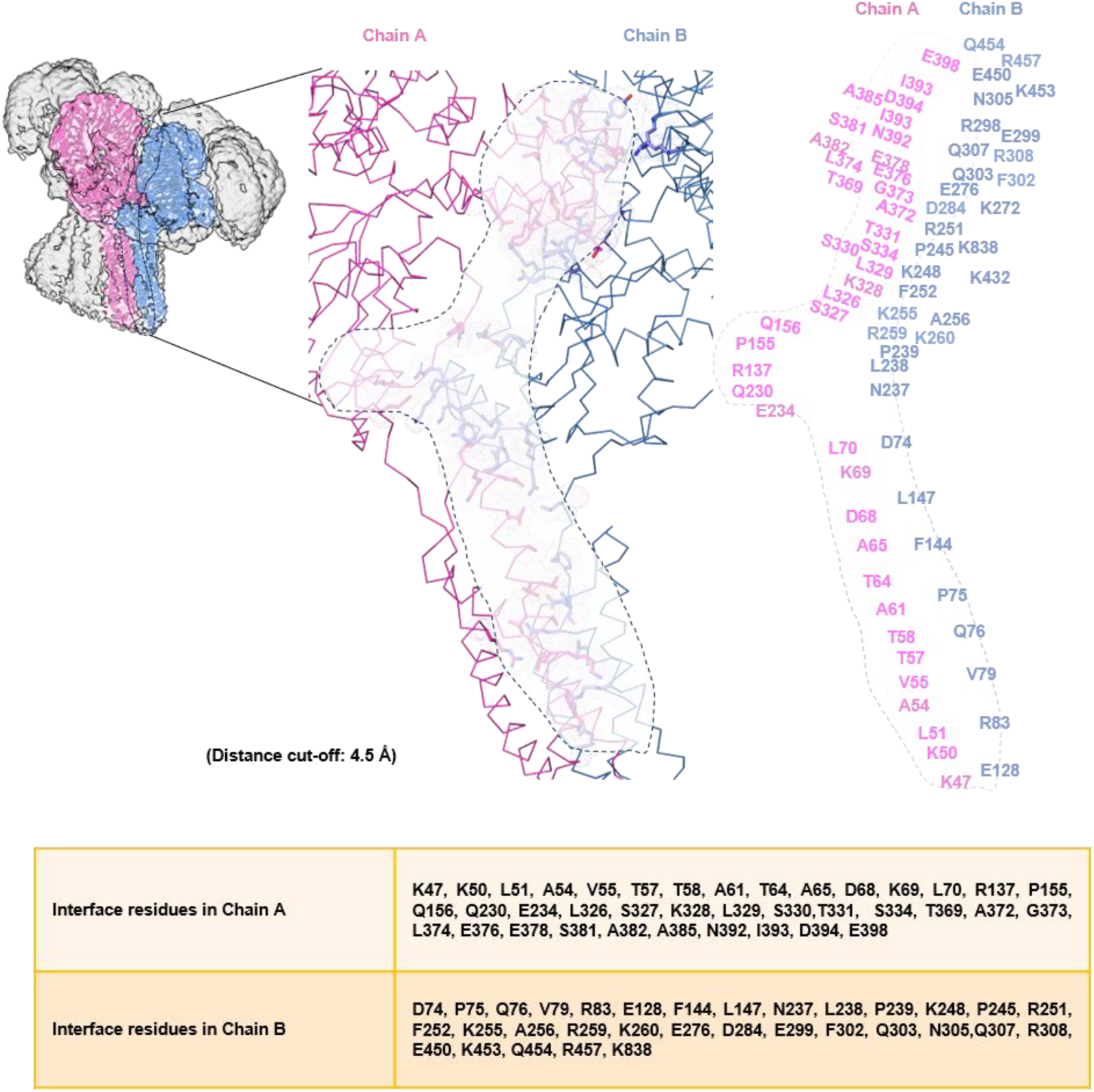
Schematic representation of interface residues holding the protomers in the WAI3 octamer. The table lists interface residues between chains A and B of the WAI3 octamer, identified using a distance cut-off of 4.5 Å.

**Figure S5.**
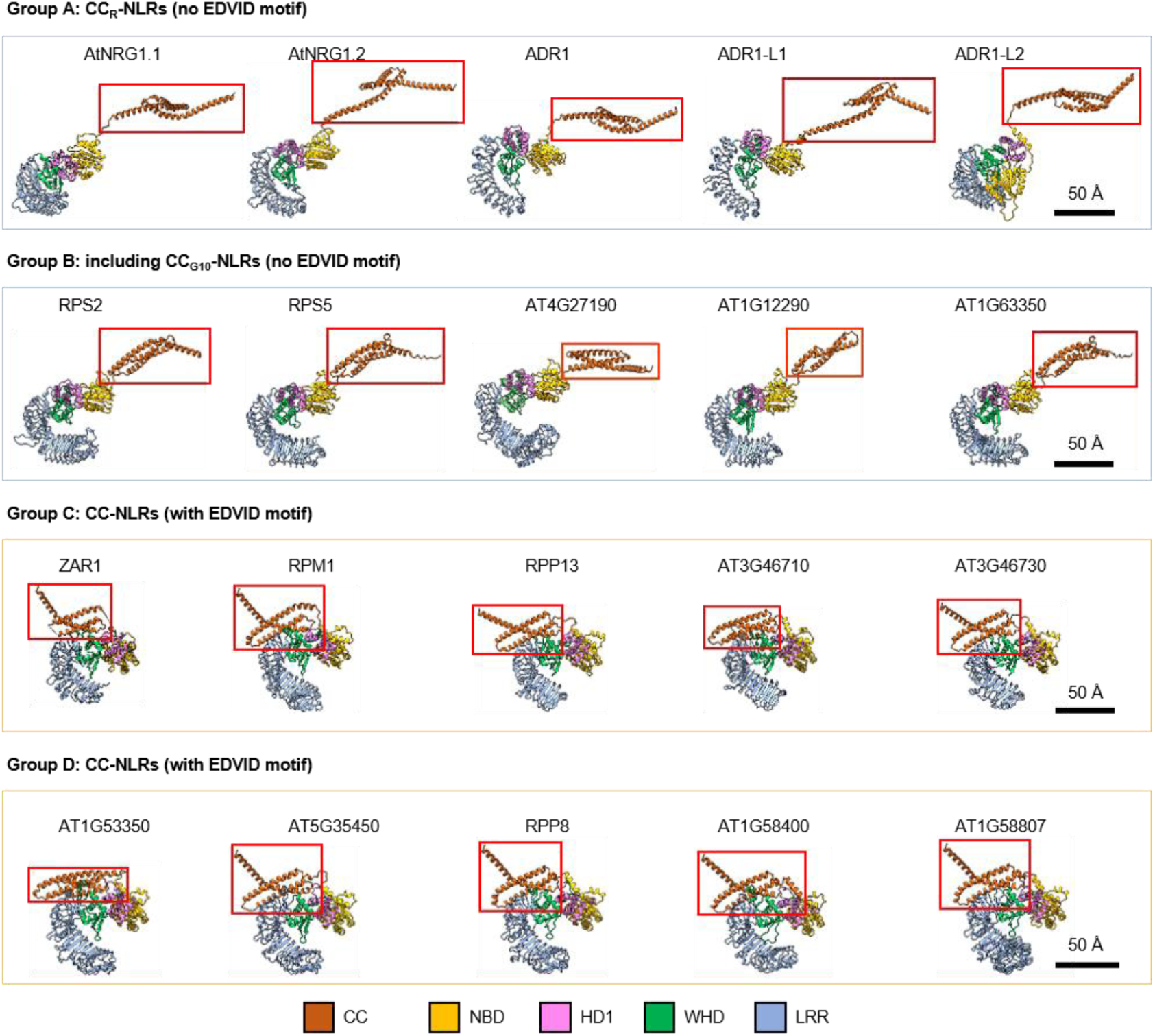
*In silico* analysis reveals a strong correlation between CC domain positioning and the presence of the EDVID motif across different CC-NLR subgroups. Based on a previously published phylogenetic analysis, *Arabidopsis* CC-NLRs can be grouped into four groups according to their CC domain sequences: Group A (e.g., helper NLRs such as NRG1 and ADR1), Group B (including CC_G10_-NLRs), and Groups C and D (CC-NLRs including ZAR1 and PRR8). Groups A and B lack the EDVID motif, whereas Groups C and D consistently contain it (Wroblewski et al., 2018). For each group, both functionally characterized and randomly selected NLRs were structurally modelled using AlphaFold3 (Abramson et al., 2024). The predictions reveal a strong correlation: CC-NLRs containing the EDVID motif adopt a ZAR1-like configuration, while those lacking the motif exhibit a WAI3-like configuration. The ZAR1 structure was adapted from the resolved structure (PDB: 6J5T), while all other models were generated using AlphaFold3 (Abramson et al., 2024). CC domains are highlighted in red boxes.

**Figure S6.**
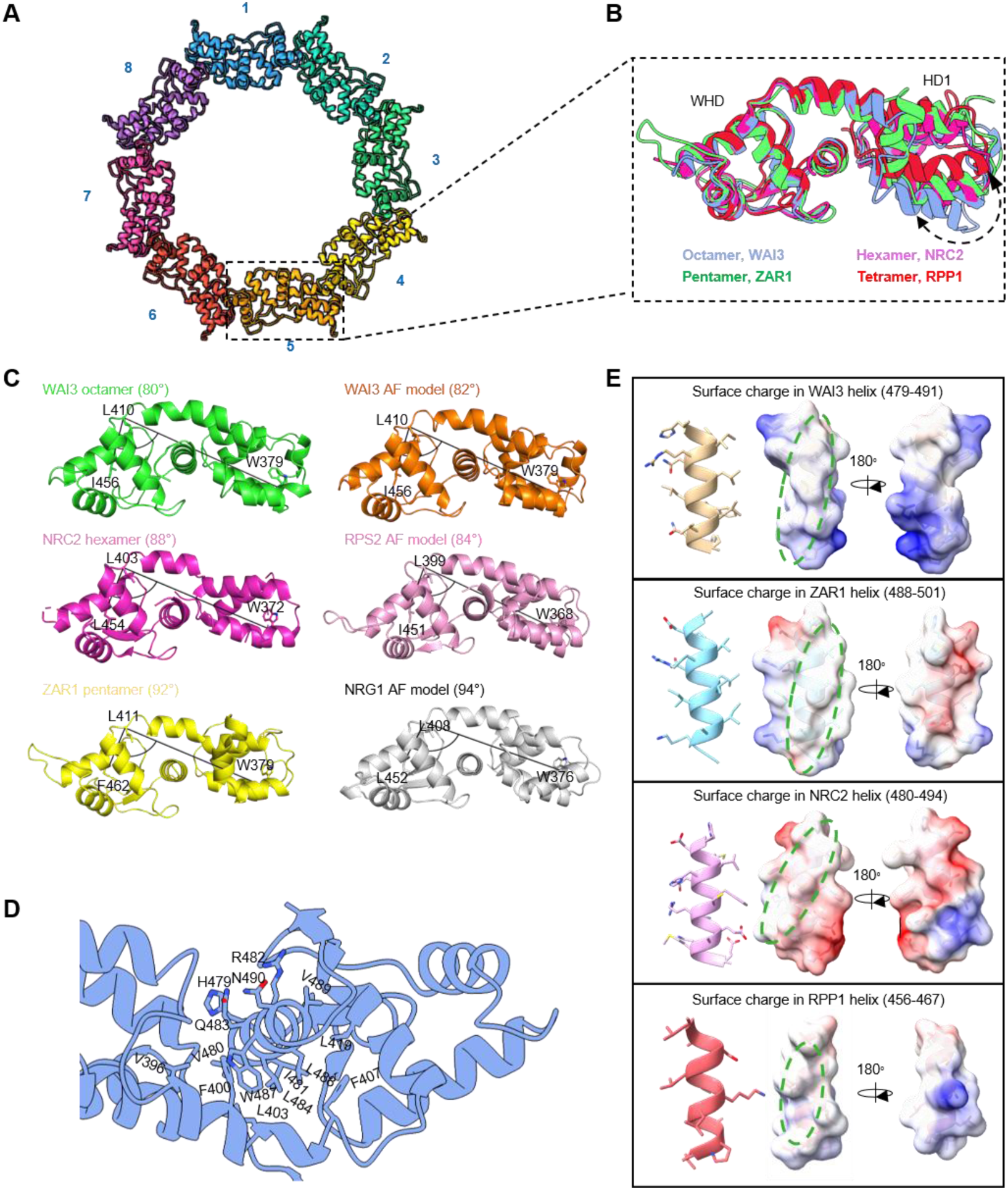
Conformational dynamics of the HD1–WHD module in resistosomes with different oligomeric states. (A) In the WAI3 octameric resistosome, the HD1–WHD domains from individual protomers form the peripheral rim of the complex. (B) Structural superposition of HD1–WHD domains from resistosomes of varying oligomeric states. (C) Quantification of HD1–WHD domain orientation in published cryo-EM structures of plant resistosomes and selected AlphaFold models. The angle between HD1 and WHD relative to the central helix was measured to assess conformational variation. (D) Structural view of the amphipathic helix (residues 479–493) and its associated hydrophobic residues. (E) Surface electrostatic potential of the amphipathic helix in WAI3, ZAR1, NRC2, and RPP1, highlighting differences in surface charge distribution.

## REFERENCES

Abramson, J., Adler, J., Dunger, J., Evans, R., Green, T., Pritzel, A., Ronneberger, O., Willmore, L., Ballard, A.J., Bambrick, J., et al. (2024). Accurate structure prediction of biomolecular interactions with AlphaFold 3. Nature 630, 493–500.

Afonine, P.V., Poon, B.K., Read, R.J., Sobolev, O.V., Terwilliger, T.C., Urzhumtsev, A., and Adams, P.D. (2018). Real-space refinement in PHENIX for cryo-EM and crystallography. Acta Crystallographica Section D 74, 531–544.

Afzal, A.J., Alam, M., Huang, J., Agha, M.A., da Cunha, L., Iqbal Rai, M., Tahir, J., Jamroze, A., Jones, J.D.G., and Mackey, D. (2025). Defense-suppressive fragments of RIN4 generated by AvrRpt2 participate in NDR1-dependent activation of RPS2. bioRxiv, 2025.2006.2002.656948.

Bi, G., Su, M., Li, N., Liang, Y., Dang, S., Xu, J., Hu, M., Wang, J., Zou, M., Deng, Y., et al. (2021). The ZAR1 resistosome is a calcium-permeable channel triggering plant immune signaling. Cell 184, 3528–3541.e3512.

Burdett, H., Bentham, A.R., Williams, S.J., Dodds, P.N., Anderson, P.A., Banfield, M.J., and Kobe, B. (2019). The Plant “Resistosome”: Structural insights into immune signaling. Cell Host Microbe 26, 193–201.

Charpentier, M., Sun, J., Martins, T.V., Radhakrishnan, G.V., Findlay, K., Soumpourou, E., Thouin, J., Véry, A.-A., Sanders, D., Morris, R.J., et al. (2016). Nuclear-localized cyclic nucleotide–gated channels mediate symbiotic calcium oscillations. Science 352, 1102–1105.

Cheng, K., Wilkinson, M., Chaban, Y., and Wigley, D.B. (2020). A conformational switch in response to Chi converts RecBCD from phage destruction to DNA repair. Nature Structural & Molecular Biology 27, 71–77.

Contreras, M.P., Lüdke, D., Pai, H., Toghani, A., and Kamoun, S. (2023). NLR receptors in plant immunity: making sense of the alphabet soup. EMBO reports 24.

Croll, T.I. (2018). ISOLDE: a physically realistic environment for model building into low-resolution electron-density maps. Acta Crystallogr D Struct Biol 74, 519–530.

Day, B., Dahlbeck, D., Huang, J., Chisholm, S.T., Li, D., and Staskawicz, B.J. (2005). Molecular basis for the RIN4 negative regulation of RPS2 disease resistance. Plant Cell 17, 1292–1305.

DeFalco, T.A., Toyota, M., Phan, V., Karia, P., Moeder, W., Gilroy, S., and Yoshioka, K. (2017). Using GCaMP3 to study Ca^2+^ signaling in *Nicotiana* species. Plant and Cell Physiology 58, 1173–1184.

Emsley, P., and Cowtan, K. (2004). Coot: model-building tools for molecular graphics. Acta Crystallogr D Biol Crystallogr 60, 2126–2132.

Engler, C., Youles, M., Gruetzner, R., Ehnert, T.M., Werner, S., Jones, J.D., Patron, N.J., and Marillonnet, S. (2014). A golden gate modular cloning toolbox for plants. ACS Synth Biol 3, 839–843.

Förderer, A., Li, E., Lawson, A.W., Deng, Y.-n., Sun, Y., Logemann, E., Zhang, X., Wen, J., Han, Z., Chang, J., et al. (2022). A wheat resistosome defines common principles of immune receptor channels. Nature 610, 532–539.

Horsefield, S., Burdett, H., Zhang, X., Manik, M.K., Shi, Y., Chen, J., Qi, T., Gilley, J., Lai, J.-S., Rank, M.X., et al. (2019). NAD^+^ cleavage activity by animal and plant TIR domains in cell death pathways. Science 365, 793–799.

Huang, S., Jia, A., Ma, S., Sun, Y., Chang, X., Han, Z., and Chai, J. (2023). NLR signaling in plants: from resistosomes to second messengers. Trends Biochem Sci 48, 776–787.

Huang, S., Wang, J., Song, R., Jia, A., Xiao, Y., Sun, Y., Wang, L., Mahr, D., Wu, Z., Han, Z., et al. (2025). Balanced plant helper NLR activation by a modified host protein complex. Nature 639, 447–455.

Jacob, P., Kim Nak, H., Wu, F., El-Kasmi, F., Chi, Y., Walton William, G., Furzer Oliver, J., Lietzan Adam, D., Sunil, S., Kempthorn, K., et al. (2021). Plant “helper” immune receptors are Ca^2+^-permeable nonselective cation channels. Science 373, 420–425.

Jia, A., Huang, S., Song, W., Wang, J., Meng, Y., Sun, Y., Xu, L., Laessle, H., Jirschitzka, J., Hou, J., et al. (2022). TIR-catalyzed ADP-ribosylation reactions produce signaling molecules for plant immunity. Science 377, eabq8180.

Jones, J.D., and Dangl, J.L. (2006). The plant immune system. Nature 444, 323–329.

Jones, J.D., Vance, R.E., and Dangl, J.L. (2016). Intracellular innate immune surveillance devices in plants and animals. Science 354.

Jones, J.D.G., Staskawicz, B.J., and Dangl, J.L. (2024). The plant immune system: From discovery to deployment. Cell 187, 2095–2116.

Jubic, L.M., Saile, S., Furzer, O.J., El Kasmi, F., and Dangl, J.L. (2019). Help wanted: helper NLRs and plant immune responses. Curr Opin Plant Biol 50, 82–94.

Kimanius, D., Dong, L., Sharov, G., Nakane, T., and Scheres, S.H.W. (2021). New tools for automated cryo-EM single-particle analysis in RELION-4.0. Biochemical Journal 478, 4169–4185.

Kourelis, J., Sakai, T., Adachi, H., and Kamoun, S. (2021). RefPlantNLR is a comprehensive collection of experimentally validated plant disease resistance proteins from the NLR family. PLOS Biology 19.

Kumar, S., Stecher, G., Li, M., Knyaz, C., Tamura, K., and Battistuzzi, F.U. (2018). MEGA X: Molecular Evolutionary Genetics Analysis across Computing Platforms. Molecular Biology and Evolution 35, 1547–1549.

Lee, H.Y., Mang, H., Choi, E., Seo, Y.E., Kim, M.S., Oh, S., Kim, S.B., and Choi, D. (2021). Genome-wide functional analysis of hot pepper immune receptors reveals an autonomous NLR clade in seed plants. New Phytol 229, 532–547.

Letunic, I., and Bork, P. (2024). Interactive Tree of Life (iTOL) v6: recent updates to the phylogenetic tree display and annotation tool. Nucleic Acids Research 52, W78–W82.

Liu, F., Yang, Z., Wang, C., You, Z., Martin, R., Qiao, W., Huang, J., Jacob, P., Dangl, J.L., Carette, J.E., et al. (2024). Activation of the helper NRC4 immune receptor forms a hexameric resistosome. Cell 187, 4877–4889 e4815.

Ma, S., Lapin, D., Liu, L., Sun, Y., Song, W., Zhang, X., Logemann, E., Yu, D., Wang, J., Jirschitzka, J., et al. (2020). Direct pathogen-induced assembly of an NLR immune receptor complex to form a holoenzyme. Science 370, eabe3069.

Madhuprakash, J., Toghani, A., Contreras, M.P., Posbeyikian, A., Richardson, J., Kourelis, J., Bozkurt, T.O., Webster, M.W., and Kamoun, S. (2024). A disease resistance protein triggers oligomerization of its NLR helper into a hexameric resistosome to mediate innate immunity. Science Advances 10, eadr2594.

Manik, M.K., Shi, Y., Li, S., Zaydman, M.A., Damaraju, N., Eastman, S., Smith, T.G., Gu, W., Masic, V., Mosaiab, T., et al. (2022). Cyclic ADP ribose isomers: Production, chemical structures, and immune signaling. Science 377, eadc8969.

Martin, R., Qi, T., Zhang, H., Liu, F., King, M., Toth, C., Nogales, E., and Staskawicz, B.J. (2020). Structure of the activated ROQ1 resistosome directly recognizing the pathogen effector XopQ. Science 370, eabd9993.

Pettersen, E.F., Goddard, T.D., Huang, C.C., Meng, E.C., Couch, G.S., Croll, T.I., Morris, J.H., and Ferrin, T.E. (2021). UCSF ChimeraX: Structure visualization for researchers, educators, and developers. Protein Science 30, 70–82.

Punjani, A., Rubinstein, J.L., Fleet, D.J., and Brubaker, M.A. (2017). cryoSPARC: algorithms for rapid unsupervised cryo-EM structure determination. Nature Methods 14, 290–296.

Rohou, A., and Grigorieff, N. (2015). CTFFIND4: Fast and accurate defocus estimation from electron micrographs. Journal of Structural Biology 192, 216–221.

Schrödinger, L., & DeLano, W. (2020). PyMOL. Retrieved from http://www.pymol.org/pymol.

Seiferth, D., and Biggin, P.C. (2024). Exploring the influence of pore shape on conductance and permeation. Biophys J 123, 3107–3119.

Selvaraj, M., Toghani, A., Pai, H., Sugihara, Y., Kourelis, J., Yuen, E.L.H., Ibrahim, T., Zhao, H., Xie, R., Maqbool, A., et al. (2024). Activation of plant immunity through conversion of a helper NLR homodimer into a resistosome. PLoS Biol 22, e3002868.

Seo, E., Kim, S., Yeom, S.I., and Choi, D. (2016). Genome-wide comparative analyses reveal the dynamic evolution of nucleotide-binding leucine-rich repeat gene family among *Solanaceae* plants. Front Plant Sci 7, 1205.

Sulkowski, O., Ovodova, A., Leisse, A., Gögelein, K., and Förderer, A. (2025). Diversification of the “EDVID” packing motif underpins structural and functional variation in plant NLR coiled-coil domains. bioRxiv, 2025.2006.2001.657260.

Wan, L., Essuman, K., Anderson, R.G., Sasaki, Y., Monteiro, F., Chung, E.-H., Osborne Nishimura, E., DiAntonio, A., Milbrandt, J., Dangl, J.L., et al. (2019). TIR domains of plant immune receptors are NAD^+^-cleaving enzymes that promote cell death. Science 365, 799–803.

Wang, J., Hu, M., Wang, J., Qi, J., Han, Z., Wang, G., Qi, Y., Wang, H.W., Zhou, J.M., and Chai, J. (2019). Reconstitution and structure of a plant NLR resistosome conferring immunity. Science 364.

Wöhlert, D., Grötzinger, M.J., Kühlbrandt, W., and Yildiz, Ö. (2015). Mechanism of Na^+^-dependent citrate transport from the structure of an asymmetrical CitS dimer. eLife 4, e09375.

Wroblewski, T., Spiridon, L., Martin, E.C., Petrescu, A.J., Cavanaugh, K., Truco, M.J., Xu, H., Gozdowski, D., Pawlowski, K., Michelmore, R.W., et al. (2018). Genome-wide functional analyses of plant coiled-coil NLR-type pathogen receptors reveal essential roles of their N-terminal domain in oligomerization, networking, and immunity. PLoS Biol 16, e2005821.

Wu, Y., Xu, W., Zhao, G., Lei, Z., Li, K., Liu, J., Huang, S., Wang, J., Zhong, X., Yin, X., et al. A canonical protein complex controls immune homeostasis and multipathogen resistance. Science 0, eadr2138.

Xiao, Y., Wu, X., Wang, Z., Ji, K., Zhao, Y., Zhang, Y., and Wan, L. (2025). Activation and inhibition mechanisms of a plant helper NLR. Nature 639, 438–446.

Xie, J., Guo, G., Wang, Y., Hu, T., Wang, L., Li, J., Qiu, D., Li, Y., Wu, Q., Lu, P., et al. (2020). A rare single nucleotide variant in *Pm5e* confers powdery mildew resistance in common wheat. New Phytol 228, 1011–1026.

Yu, D., Song, W., Tan, E.Y.J., Liu, L., Cao, Y., Jirschitzka, J., Li, E., Logemann, E., Xu, C., Huang, S., et al. (2022). TIR domains of plant immune receptors are 2′,3′-cAMP/cGMP synthetases mediating cell death. Cell 185, 2370–2386.e2318.

Yu, H., Xu, W., Chen, S., Wu, X., Rao, W., Liu, X., Xu, X., Chen, J., Nishimura, M.T., Zhang, Y., et al. Activation of a helper NLR by plant and bacterial TIR immune signaling. Science 0, eadr3150.

Zhao, Y.-B., Liu, M.-X., Chen, T.-T., Ma, X., Li, Z.-K., Zheng, Z., Zheng, S.-R., Chen, L., Li, Y.-Z., Tang, L.-R., et al. (2022). Pathogen effector AvrSr35 triggers Sr35 resistosome assembly via a direct recognition mechanism. Science Advances 8, eabq5108.

Zheng, S.Q., Palovcak, E., Armache, J.-P., Verba, K.A., Cheng, Y., and Agard, D.A. (2017). MotionCor2: anisotropic correction of beam-induced motion for improved cryo-electron microscopy. Nature Methods 14, 331–332.

